# Isolating the Sources of Pipeline-Variability in Group-Level Task-fMRI results

**DOI:** 10.1101/2021.07.27.453994

**Authors:** Alexander Bowring, Thomas E. Nichols, Camille Maumet

## Abstract

While the development of tools and techniques has broadened our horizons for comprehending the complexities of the human brain, a growing body of research has highlighted the pitfalls of such methodological plurality. In a recent study, we found that the choice of software package used to run the analysis pipeline can have a considerable impact on the final group-level results of a task-fMRI investigation (Bowring et al., 2019, *BMN*). Here we revisit our work, seeking to identify the stages of the pipeline where the greatest variation between analysis software is induced. We carry out further analyses on the three datasets evaluated in *BMN*, employing a common processing strategy across parts of the analysis workflow and then utilizing procedures from three software packages (AFNI, FSL and SPM) across the remaining steps of the pipeline. We use quantitative methods to compare the statistical maps and isolate the main stages of the workflow where the three packages diverge. Across all datasets, we find that variation between the packages’ results is largely attributable to a handful of individual analysis stages, and that these sources of variability were heterogeneous across the datasets (e.g. choice of first-level signal model had the most impact for the ds000001 dataset, while first-level noise model was more influential for ds000109 dataset). We also observe areas of the analysis workflow where changing the software package causes minimal differences in the final results, finding that the group-level results were largely unaffected by which software package is used to model the low-frequency fMRI drifts.

## 1. Introduction

Public speaking could be the most nerve-jangling aspect of working as a scientist. While presenting research findings is, in theory, a perfect opportunity to showcase the fruits of our labor to fellow experts in our subject area, a big talk can often be an anxiety-ridden experience characterized by sweaty palms, a dry mouth, and butterflies in the stomach. Fortunately for academics, the social psychology literature provides us with some ‘life-hacks’ for dealing with such high-stress situations. One study, published in 2010, suggested that assuming a ‘power pose’ – standing tall with an expansive, self-assured body posture – can evoke real feelings of courage and assertiveness within ourselves (Carney et al., 2010). As well as this, a famous investigation carried out in the 1980’s found that our own facial expressions can influence our psychological state of mind (Strack et al., 1988). For public speaking, this arms us with a plan of attack: by hiding any initial feelings of dread with a big smile and a ‘Wonder Woman’-like posture, academics can exert the genuine confidence and joy they wish for in their presentations!

The bad news, however, is that power-posing and faux-smiling are both examples of psychological effects that have recently been challenged. In 2016, the efforts of 17 independent teams to replicate the findings of Strack et al. (1988) all led to null-results, bringing into question the validity of the original study (Wagenmakers et al., 2016). Similarly, various attempts to replicate the effects of power-posing have also been unsuccessful (Ranehill et al., 2015, Garrison et al., 2016, Smith and Apicella, 2017), prompting a statement from one of the original study authors to retract their findings (Carney, 2017). Regrettably for academics, there may not be any shortcuts to mastering an oral presentation after all.

These aren’t the only examples of research that have recently been undermined. In fact, the 2010’s may be best remembered by social scientists as the start of the ‘replication crisis’ (Maxwell et al., 2015), an ongoing issue that has gained prominence as a number of classic and contemporary psychology studies have been brought into question. At the heart of the controversy is a growing body of work where attempts to replicate several effects in psychological science have failed, prompting further scrutiny of the robustness of the original findings. In a landmark investigation, the Open Science Collaboration (2015) repeated 100 experiments that had been published in three high-ranking psychology journals, reporting that only 36% of their replications determined a positive result compared to 97% of the original studies. At around the same time the first in a series of Many Labs studies was published, where numerous analysis teams have tried to replicate results from the psychology literature across a diverse range of samples. Of the 51 studies re-evaluated in the first three Many Labs projects, roughly 60% yielded significant effects (Klein et al., 2014, 2018, Ebersole et al., 2016).

The field of functional magnetic resonance imaging (fMRI) for human brain mapping has not come away unharmed from the replication crisis. On the contrary, the large degree of flexibility in neuroimaging analysis workflows has been pinpointed as an aspect of the field that can hinder reproducibility (Ioannidis, 2005). The crux of the problem is that two different analysis pipelines applied to the same dataset are unlikely to give the same result. Therefore, as an increasing number of analytical tools and techniques have become available to researchers, this has also increased the potential to yield distorted findings with inflated levels of false activations. When combined with selective reporting practices – where only methods that return a favourable outcome are likely to end up being published – the consequences of this can be severe, leading to fMRI effect sizes that are misrepresented and often overstated in the neuroimaging literature (Simmons et al., 2011, Szucs and Ioannidis, 2017).

In one of the most comprehensive studies in this area, a single publicly available fMRI dataset was analyzed using over 6,000 unique simulated workflows, constructed by enumerating all possible pipeline combinations from an array of commonly implemented analysis procedures (Carp, 2012). Across the tens of thousands of thresholded results maps generated by these workflows, a substantial degree of variability was observed in both the sizes and locations of significant activation. In a more recent study, 70 independent research teams were tasked with testing 9 hypotheses on the same fMRI dataset, with no constraints placed on how each team approached their analysis (Botvinik-Nezer et al., 2020). Consequently, no two teams chose the same analysis workflow, and once again the plurality of methodological approaches manifested as variability in the final scientific outcomes, this time with considerable disagreement between the 70 teams’ hypothesis test results. Overall, these investigations have forewarned practitioners not to fall victim to a version of insanity where we apply *different* workflows over and over again and expect *the same* results.

In Bowring, Maumet, and Nichols (2019) (*BMN*), we discovered that it is not just the procedures comprising the analysis pipeline that can induce variation across fMRI results, but also the choice of *software package* through which the analysis is conducted. We reanalyzed three datasets connected to three published task-fMRI studies within the three most widely-used neuroimaging software packages – AFNI, FSL, and SPM – reproducing the original publication’s analysis workflows in each package as closely as possible so that the difference in software was the only changing variable. We then applied a range of similarity metrics to quantify the differences between each software’s final group-level results. While qualitatively certain patterns of signal were observed across all three packages’ statistical results maps, our quantitative comparisons displayed marked differences in the size, magnitude and topology of activated brain regions, and we ultimately concluded that weak effects may not generalize across software.

Now we revisit that work, seeking to understand *where* in the analysis pipeline the greatest variation between analysis software is induced. We substantially extend the analyses carried out for *BMN*, running the same three datasets through a series of ‘hybrid’ pipelines that employ a common processing strategy across parts of the workflow (e.g. by implementing a common fMRIPrep preprocessing strategy) and then interchange pipeline elements between software for the remaining stages of the analysis. By comparing all sets of our analysis results,we isolate the key stages of the workflow where the three packages diverge. Ultimately, we find that the variation between the packages’ results is largely attributable to sizable processing differences at a handful of key analysis stages, but that these sources of variability can be heterogeneous across datasets. Finally, for each study we apply an image-based meta-analysis procedure recently used in Botvinik-Nezer et al. (2020) to all of our analysis results, aggregating the information acquired from running one dataset through multiple pipelines to obtain a consensus map of activated brain regions.

The remainder of the manuscript is organised as follows: First, we provide a brief summary of the three original published studies from which we sourced our selected datasets. We then describe the pipelines implemented for our reanalyses of the data, and detail the quantitative and qualitative metrics and image-based meta-analysis procedure applied to our analysis results. Finally, we evaluate our findings to assess the magnitude of variation between fMRI analysis software at each stage of the analysis workflow, and discuss the repercussions of these results on the functional neuroimaging literature.

## 2. Methods

We first provide an overview of the original study paradigms for the three published task-fMRI works from which we sourced the three datasets, before we go on to detail the reanalysis methods carried out in this work. Most notably, while the original studies’ analyses were carried out on 16, 29 and 30 participants task-fMRI data respectively, for the latter two studies only 21 and 17 participants’ data were available for reanalyses. Alongside this, due to preprocessing failing for one individual in the ds000001 dataset, we ultimately reanalyzed 15 subjects rather than the complete sample of 16 whose data were shared (see the start of Section 3 for more details).

### 2.1. Study Description and Data Source

We selected three task-fMRI studies from the publicly accessible OpenfMRI (now upgraded to OpenNeuro, RRID:SCR_005031) data repository (Gorgolewski et al., 2017), OpenfMRI dataset accession numbers: ds000001 (Revision: 2.0.4; Schonberg et al., 2012), ds000109 (Revision 2.0.2; Moran et al., 2012), and ds000120 (Revision 1.0.0; Padmanabhan et al., 2011). Each of the datasets had been organized in compliance with the Brain Imaging Data Structure (BIDS, RRID:SCR_016124; Gorgolewski et al., 2016). These datasets were chosen following an extensive selection procedure (carried out between May 2016 and November 2016), whereby we vetted the associated publication for each dataset stored in the repository. We sought to find studies with simple analysis pipelines and clearly reported regions of brain activation that would be easily comparable to our own results. Exclusion criteria included the use of custom software, activations defined using small volume correction, and application of more intricate methods such as region of interest and robust regression analysis, which we believed could be impractical to implement across all analysis software. A full description of the paradigm for each of our chosen studies is included in the respective publication, here we give a brief overview.

For the ds000001 study, 16 healthy adult subjects participated in a balloon analog risk task over three scanning sessions. On each trial, subjects were presented with a simulated balloon, and offered a monetary reward to ‘pump’ the balloon. With each successive pump the money would accumulate, and at each stage of the trial subjects had a choice of whether they wished to pump again or cash-out. After a certain number of pumps, which varied between trials, the balloon exploded. If subjects had cashed-out before this point they were rewarded with all the money they had earned during the trial, however if the balloon exploded all money accumulated was lost. Three different coloured ‘reward’ balloons were used between trials, each having a different explosion probability, as well as a grey ‘control’ balloon, which had no monetary value and would disappear from the screen after a predetermined number of pumps. Here we reproduce the pipeline used to obtain the main study result contrasting the parametrically modulated activations of pumps of the reward balloons versus pumps of the control balloon, corresponding to Figure 3 and Table 2 in the original paper.

The ds000109 study investigated the ability of people from different age-groups to understand the mental state of others. A total of 48 subjects participated, although imaging data was obtained from only 43 participants for the false belief task: 29 younger adults and 14 older adults. In this task participants listened to either a ‘false belief’ or ‘false photo’ story. A false belief story would entail an object being moved from one place to another, with certain characters witnessing the change in location while others were unaware. False photo stories were similar except that they involved some physical representation of the missing object, such as a photo of an object in a location from which it had been subsequently removed. The task had a block design where stories were represented for ten seconds, after which participants had to answer a question about one of the characters’ perceptions of the location of the object. We reproduce the pipeline used to obtain the contrast map of false belief versus false photo activations for the younger adults, corresponding to Figure 5a and Table 3 from the original publication.

Finally, the ds000120 study explored reward processing across different age groups. fMRI results were reported on 30 subjects, with 10 participants belonging to each of the three age groups (children, adolescents and adults). Participants took part in an antisaccade task where a visual stimulus was presented in each trial and subjects were instructed to quickly fixate their gaze on the side of the screen opposite to the stimulus. Prior to a trial, subjects were given a visual cue to signal whether or not they had the potential to win a monetary reward based on their upcoming performance (a ‘reward’ or ‘neutral’ trial). We reproduce the pipeline used to obtain the main effect of time activation map, an *F*-statistic for any non-zero coefficients in the sine HRF basis, corresponding to Figure 3 and Table 1 in the original publication.

**Table 1:** fMRIPrep Processing Pipeline. For all three studies, a subset of the workflows applied the same fMRIPrep preprocessing pipeline. Here, we itemize the main steps of the fMRIPrep preprocessing pipeline, making note of the various tools used at each stage of the workflow.

| Workflow | Processing Step | Description | Tools Used |
| --- | --- | --- | --- |
| **Structural Preprocessing** | Non-uniform Intensity Correction | The anatomical T1w image was corrected for intensity non-uniformity with N4BiasFieldCorrection, distributed within ANTs 2.2.0, to be used as the anatomical reference image for the rest of the pipeline. | ANTs |
|  | Brain Extraction | The anatomical reference image was skull-stipped with a Nipype implementation of the antsBrainExtraction.sh workflow from ANTs. | ANTs |
|  | Segmentation | Brain tissue segmentation of the CSF, WM and GM was performed on the brain-extracted T1w using FSL’s fast. | FSL |
|  | Brain Surface Reconstruction | Brain surfaces were reconstructed using FreeSurfer’s recon-all, after which the brain mask was refined using a custom variation of the ANTs-derived and FreeSurfer-derived segmentations of the cortical GM. | FreeSurfer, ANTs |
|  | T1w-to-MNI152 registration | Spatial normalization to MNI152 space was performed through nonlinear registration with ANTs’ antsRegistration, using brain-extracted versions of the T1w reference and template images. | ANTs |
| **Functional Preprocessing** | Reference Image | For each BOLD run, a custom methodology of fMRIPrep was applied to average across the BOLD time-series in order to generate a reference volume. | Custom |
|  | Brain Extraction | The BOLD reference image was skull-stripped using NiWorkflows’ init_enhance_and_skullstrip_bold_wf(), to be used for head-motion estimation and registration of the BOLD time-series images to the subject’s T1w image. | NiWorkflows |
|  | BOLD-to-T1w registration | The BOLD reference was co-registered to the T1w reference using FreeSurfer’s bbregister, implementing a boundary-based registration with 6 degrees of freedom. | FreeSurfer |
|  | Head-motion Correction | Head-motion parameters with respect to the BOLD reference (one rigid-body transformation, three rotations and three translations) were estimated with FSL’s mcflirt, after which the rigid-body transformation was applied to re-sample the BOLD time-series onto their original, native space. | FSL |
|  | BOLD-to-MNI152 registration | Transformations already obtained (head-motion rigid-body transformation, BOLD to T1w registration, T1w to MNI152 transformation) were concatenated to map the BOLD image to the MNI152 standard space. |  |
|  | Confound Estimation | A range of potential confounds were estimated, including the mean global signal, mean tissue class signal, tCompCor, aCompCor, Framewise Displacement and DVARS. | CompCor |

### 2.2. Previous Analyses and Preprocessing Methods

In *BMN* we reanalyzed the ds000001 and ds000109 studies described in the previous section using each of the three software packages: AFNI (version AFNI_18.1.09), RRID:SCR_005927; FSL (version 5.0.10), RRID:SCR_002823; and SPM (version SPM12, v6906), RRID:SCR_007037. For ds000120, the repeated-measures design carried out for the original group-level analysis was not feasible to implement in FSL, and therefore we reanalyzed the data in AFNI and SPM only. In parallel to our reproductions of the original analysis workflows, for ds000001 and ds000109 we computed an additional set of group-level results using the nonparametric (permutation test) inference procedures available within the three software packages. For ds000120 a one-sample repeated-measure permutation test was not viable in AFNI, so nonparametric inference was excluded for this study.

The pipelines carried out for each study and software package are described in Section 2.2 of *BMN*, and a full decomposition of the modules used within each package is provided in Table 1 of the manuscript. Notably, we chose to implement a number of processing steps for all of our reanalyses regardless of whether they had carried out in the original studies. These were procedures that we believed were fundamental to ensure our reproductions could be compared objectively, and steps that are widely considered as good practice within the community. Specifically, in all of our reanalyses we applied skull stripping to the T1-weighted (T1w) structural image, we used each package’s nonlinear registration tools to transform the structural and functional data to the anatomical template, and 6 motion regressors were included in the analysis design matrix for all pipelines (while more than 6 motion regressors are often used, we chose 6 as this could be easily implemented in all three software packages).

Here, our first aim was to isolate whether the largest variation between software occurs during the preprocessing or statistical modelling of the functional data. To do this, we repeated the *BMN* analyses, except this time implementing *the same* preprocessing strategy to the three datasets before carrying out the rest of the analyses in the three packages. A common minimal preprocessing workflow was applied to each of the datasets using fMRIPrep 20.0.02 (Esteban et al., 2019, 2020b; RRID:SCR_016216), which is based on Nipype 1.4.2 (Gorgolewski et al., 2011, Esteban et al., 2020a; RRID:SCR_002502). The fMRIPrep pipeline combines procedures from a range of software packages to provide the optimal implementation at each stage of preprocessing. We will now describe the preprocessing sub-workflows that were applied to all three datasets’ anatomical and functional data within fMRIPrep. These pipelines are also summarised in Table 1, where we have included the tools implemented by fMRIPrep at each processing step. Notably, apart from a few procedures that relied on tools from FSL, most of the preprocessing performed by fMRIPrep used packages independent of AFNI, FSL and SPM.

#### 2.2.1. Anatomical Data Preprocessing in fMRIPrep

For each of the three datasets, the preprocessing of the anatomical data was carried out within fMRIPrep as follows. The T1w image was corrected for intensity non-uniformity (INU) with N4BiasFieldCorrection (Tustison et al., 2010), distributed with ANTs 2.2.0 (Avants et al., 2008, RRID:SCR_004757), and used as T1w-reference throughout the workflow. The T1w-reference was then skull-stripped with a Nipype implementation of the antsBrainExtraction.sh workflow (from ANTs), using OASIS30ANTs as a target template. Brain tissue segmentation of cerebrospinal fluid (CSF), white-matter (WM) and gray-matter (GM) was performed on the brain-extracted T1w using fast (FSL 5.0.9, RRID:SCR_002823, Zhang et al., 2001). Brain surfaces were reconstructed using recon-all (FreeSurfer 6.0.1, RRID:SCR_001847, Dale et al., 1999), and the brain mask estimated previously was refined with a custom variation of the method to reconcile ANTs-derived and FreeSurfer-derived segmentations of the cortical gray-matter of Mindboggle (RRID:SCR_002438, Klein et al., 2017). Volume-based spatial normalization to one standard space (MNI152NLin2009cAsym) was performed through nonlinear registration with antsRegistration (ANTs 2.2.0), using brain-extracted versions of both T1w reference and the T1w template. The following template was selected for spatial normalization: ICBM 152 Nonlinear Asymmetrical template version 2009c [Fonov et al., 2009, RRID:SCR_008796; TemplateFlow ID: MNI152NLin2009cAsym].

#### 2.2.2. Functional Data Preprocessing in fMRIPrep

For each of the the three datasets, the preprocessing of the functional data was carried out within fMRIPrep as follows. For each of the BOLD runs found per subject (across all tasks and sessions), the following preprocessing was performed. First, a reference volume and its skull-stripped version were generated using a custom methodology of fMRIPrep. Susceptibility distortion correction (SDC) was omitted. The BOLD reference was then co-registered to the T1w reference using bbregister (FreeSurfer) which implements boundary-based registration (Greve and Fischl, 2009). Co-registration was configured with six degrees of freedom.

Head-motion parameters with respect to the BOLD reference (transformation matrices, and six corresponding rotation and translation parameters) are estimated before any spatiotemporal filtering using mcflirt (FSL 5.0.9,Jenkinson et al., 2002). The BOLD time-series (including slice-timing correction when applied) were resampled onto their original, native space by applying the transforms to correct for head-motion. These resampled BOLD time-series will be referred to as preprocessed BOLD in original space, or just preprocessed BOLD. The BOLD time-series were resampled into standard space, generating a preprocessed BOLD run in MNI152NLin2009cAsym space. First, a reference volume and its skull-stripped version were generated using a custom methodology of fMRIPrep. Several confounding time-series were calculated based on the preprocessed BOLD: framewise displacement (FD), DVARS and three region-wise global signals. FD and DVARS are calculated for each functional run, both using their implementations in Nipype (following the definitions by Power et al., 2014). The three global signals are extracted within the CSF, the WM, and the whole-brain masks. Additionally, a set of physiological regressors were extracted to allow for component-based noise correction (CompCor, Behzadi et al., 2007). Principal components are estimated after high-pass filtering the preprocessed BOLD time-series (using a discrete cosine filter with 128s cut-off) for the two CompCor variants: temporal (tCompCor) and anatomical (aCompCor). tCompCor components are then calculated from the top 5% variable voxels within a mask covering the subcortical regions. This subcortical mask is obtained by heavily eroding the brain mask, which ensures it does not include cortical GM regions. For aCompCor, components are calculated within the intersection of the aforementioned mask and the union of CSF and WM masks calculated in T1w space, after their projection to the native space of each functional run (using the inverse BOLD-to-T1w transformation). Components are also calculated separately within the WM and CSF masks. For each CompCor decomposition, the k components with the largest singular values are retained, such that the retained components’ time series are sufficient to explain 50 percent of variance across the nuisance mask (CSF, WM, combined, or temporal). The remaining components are dropped from consideration. The head-motion estimates calculated in the correction step were also placed within the corresponding confounds file. The confound time series derived from head motion estimates and global signals were expanded with the inclusion of temporal derivatives and quadratic terms for each (Satterthwaite et al., 2013). Frames that exceeded a threshold of 0.5 mm FD or 1.5 standardised DVARS were annotated as motion outliers. All resamplings can be performed with a single interpolation step by composing all the pertinent transformations (i.e. head-motion transform matrices, susceptibility distortion correction when available, and co-registrations to anatomical and output spaces). Gridded (volumetric) resamplings were performed using antsApplyTransforms (ANTs), configured with Lanczos interpolation to minimize the smoothing effects of other kernels (Lanczos, 1964). Non-gridded (surface) resamplings were performed using mri_vol2surf (FreeSurfer).

### 2.3. Manipulation of Modelling Methods and Hybrid Pipeline Generation

Aside from preprocessing, different parts of the three software packages’ pipelines were interchanged to generate a collection of hybrid analysis pipelines. For these pipelines, AFNI (version AFNI_20.0.20), FSL (version 6.0.3) and SPM (standalone version of SPM12, r7771) were used. At the subject-level, modelling was partitioned into three separate components: the fMRI signal model, noise model, and low-frequency drift model; at the group-level all packages share the same one-sample signal model but are differentiated by their noise model. Notably, for the subject-level analyses it was not feasible to implement one software’s noise model inside another package (e.g. it was not viable to conduct a workflow in SPM that implemented FSL’s first-level noise model). However, exchanging the fMRI signal and low-frequency drift models between software could be done easily, by simply interchanging the relevant regressors in the design matrix. Because of this, for each of our hybrid analysis pipelines the choice of software used for modelling the noise ultimately determined the package through which the subject-level analyses were conducted. For example, a hybrid pipeline using FSL’s first-level noise model with AFNI’s first-level fMRI signal model and drift model would be implemented within FSL, except the regressors in the design matrix for modelling the fMRI signal and low-frequency drifts were then interchanged with the corresponding regressors from the design matrix generated by running the complete analysis within AFNI. In addition to this, six motion parameters (translations and rotations) estimated by fMRIPrep as part of the preprocessing workflow were included in all first-level analysis models as nuisance regressors. Finally, for each hybrid pipeline the subject-level contrast of parameter estimate maps were inputted into the software package specified by the workflow for group-level analysis and inference.

Taking all combinations of software procedures considered across the the three datasets yielded a total of 59 unique workflows, shown diagrammatically in Figure 1. The diagrams labelled **1** (far-left) and **7** (far-right) for each study display the pipelines that were carried out in *BMN*, where a single software package was used to conduct the entire analysis workflow. Pipelines labelled **2** and **6** are similar, except that preprocessing was carried out using the fMRIPrep workflow described in the previous section. Pipelines labelled **3** to **5** include further manipulations, each step interchanging one aspect of the subject- and group-level modelling as described above. For ds000001 and ds000109, modifications to the workflow were considered relative to the pipeline where the entire analysis was carried out within FSL (labelled **7F** in Fig. 1, the ‘reference’ pipeline). In other words, pipelines were generated by sequentially exchanging procedures from FSL with the corresponding procedures from AFNI (**AF** pipelines) or SPM (**SF** pipelines). For ds000120, where group-level analysis in FSL was not feasible, the pipeline carried out entirely in SPM (labelled **7S** in Fig. 1) was used as the reference instead, and pipelines were generated by exchanging procedures from SPM with the corresponding procedures from AFNI. As previously discussed, for ds000001 and ds000109 we considered both parametric and nonparametric inference (purple lines in Fig. 1) at the group-level, while for ds000120 the repeated-measures permutation test was not feasible in AFNI and therefore only parametric inference was considered.

**Figure 1:**
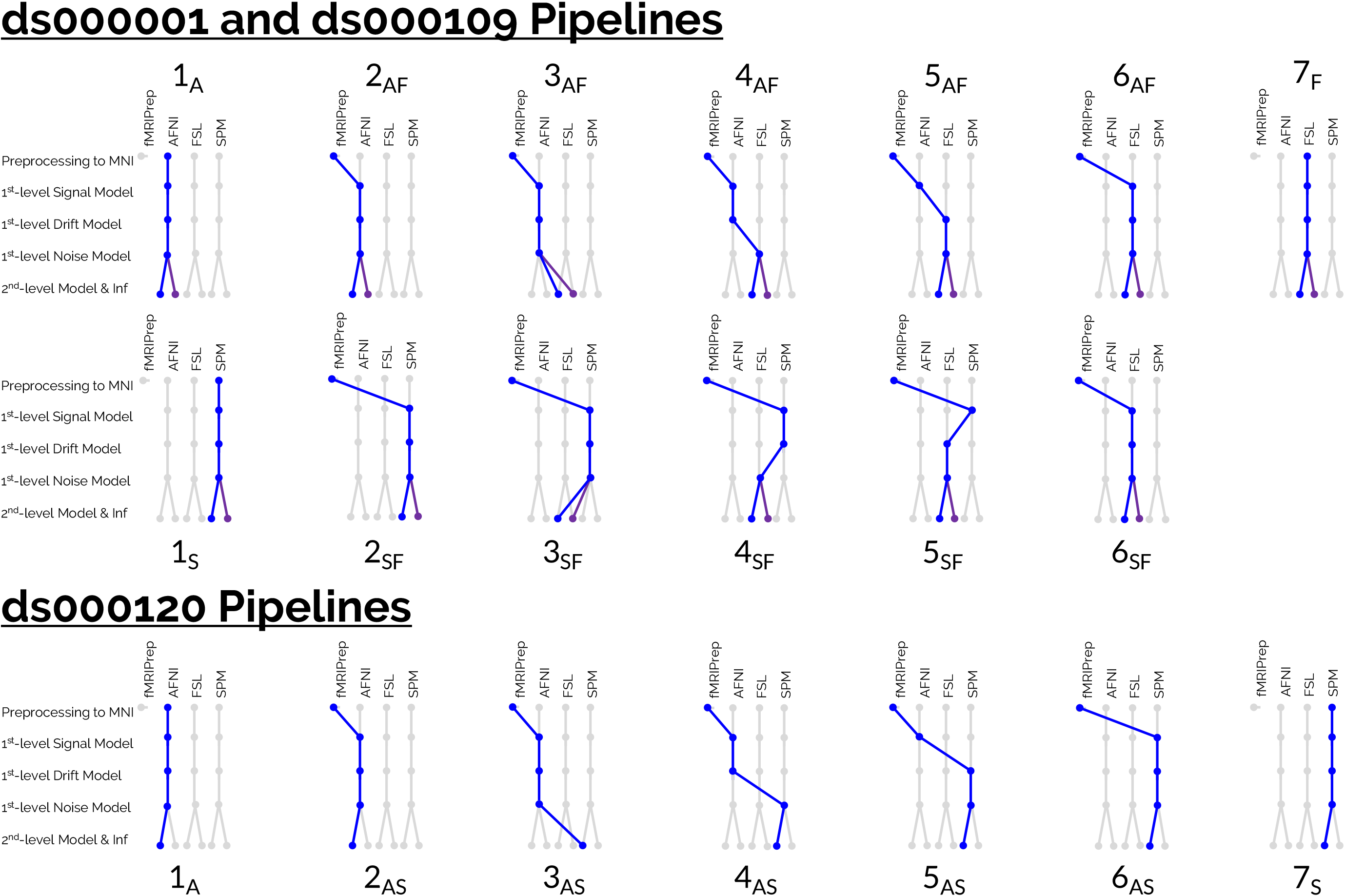
Diagrams to enumerate the complete set of 59 pipelines that were carried out on the three datasets. For ds000001 and ds000109, 26 pipelines were implemented for each dataset (13 workflows using parametric inference at the group-level, and 13 parallel workflows using nonparametric inference displayed by the purple lines). For these two datasets, FSL was used as the reference software package, and hybrid pipelines were generated by interchanging procedures from either AFNI and FSL (**AF** pipelines) or SPM and FSL (**SF** pipelines) across the analysis workflow. A total of 7 pipelines were carried out on the ds000120 dataset, where it was not feasible to analyze the data in FSL and nonparametric inference was unavailable in AFNI. Here, SPM was used as the reference software package, and hybrid pipelines were generated by interchanging procedures from AFNI and SPM across the analysis workflow. Notably, in this arrangement only one specific analysis procedure is changed between adjacent pipelines (from left-to-right). For instance, the only difference between the third and fourth pipelines on the top row was whether AFNI’s or FSL’s first-level noise model was applied, and therefore any discrepancies between the group-level results for these pipelines are wholly attributable to differences between the two softwares’ noise models.

### 2.4. Comparison Methods

Two comparison methods were considered to assess the nature of pipeline-variability across each studies’ collection of group-level statistical results. Correlations (Pearson’s *r*) were obtained for each pair of unthresholded group-level statistic maps to evaluate differences in the overall activation profiles produced from each analysis workflow. As well as this, Dice coefficients were obtained for all pairwise combinations of thresholded statistic maps in order to compare the final locations of activation given by each analysis pipeline after correction for multiple comparisons. For a pair of thresholded maps, the Dice coefficient is calculated as the volume of the intersection of the two maps divided by the average of the volume’s of each separate thresholded image. In other words, Dice measures the overlap of voxels between two sets of thresholded maps relative to the total spatial extent covered by both maps’ activations (a Dice coeffecient of 1 indicates identical locations of activation in both maps, while 0 indicates complete disagreement). With each Dice coefficient, the percentage of ‘spill-over’ activation was also computed, that is, the percentage of activation in one pipeline’s thresholded statistic map that fell outside of the analysis mask of the other pipeline.

Finally, we applied a recently proposed image-based meta-analysis method that aggregated information across all pipelines (for a given dataset) to yield a ‘consensus’ activation map, i.e. the set of brain regions where significant activation was unanimous across *all* analysis pipelines applied to the data. The consensus analysis was performed on the collection of unthresholded group-level *z*-statistic images obtained across all pipelines, accounting for the correlations between pipelines owing to the same underlying data and identical procedures applied across parts of the analysis workflow. The method was originally proposed in Botvinik-Nezer et al., 2020, where it was used to infer a consensus across results obtained by many analysis teams for a single task-fMRI dataset. Full details of the method are provided in Supplementary Section 8.1. We applied the consensus analysis methods to further examine the robustness of the individual results obtained for each dataset after accounting for the inter-pipeline variation.

For new analyses (all pipelines in Fig. 1 excluding diagrams labelled **1** and **7** that were carried out as part of *BMN*), AFNI and FSL scripts were written in Python 3.7.6 and SPM scripts were written in GNU Octave (version 4.4.1). Scripts were made generalizable using a series of templates to extract the stimulus timings from the raw data, carry out the fMRIPrep preprocessing workflow, and subsequently conduct subject- and group-level analyses. A master script for each dataset took the templates as inputs, replacing various holding variables to create distinct batch scripts for each of the unique pipelines. These batch scripts were subsequently executed within the master script to obtain all sets of group-level results.

## 3. Results

All analysis scripts and results have been made available, see Section 5 for more details.

The preprocessing of each subject’s data for all three studies was assessed using the summary reports provided as part of the fMRIPrep workflow. This included checking that each particant’s functional data had been correctly masked and successfully registered to the MNI template image. Inspection of these reports confirmed that preprocessing had been successful for all-but-one subject, subject 4 from the ds000001 dataset. Exceptionally high intensities found in this subject’s raw T1w anatomical image (potentially due to an erroneous brain-extraction applied to the anatomical data before it was shared) caused drop-out in sizable regions of the brain during bias-field correction in the fMRIPrep preprocessing pipeline. This subsequently led to a highly shrunken brain mask, and failure to register this subject’s functional data to the template image. For these reasons subject 4 was excluded from all further analyses, and we repeated all ds000001 analyses that were previously carried out for *BMN* with this subject removed. As such, all ds000001 results presented here are for 15 subjects rather than the complete sample of 16 whose data were originally shared.

### 3.1. Main Sources of Pipeline-Variability

A detailed review of the regions of activation found in the thresholded maps across all pipelines for each dataset is provided in Supplementary Section 8.2. Here we describe the main sources of variation observed between the pipelines’ results across the three datasets.

Our analyses of the ds000001 dataset suggest that differences in each software’s first-level signal model were the largest contributor to variation across the three software packages’ final statistical results. This is highlighted in Figure 2 and (Supplementary) Figure S1 (All supplementary figures described in this section can be found in Supplementary Section 8.3), where in both figures we compare the results from the two analysis workflows which differed only by the choice of first-level signal model: In Fig. 2, pipeline **5SF** applied SPM’s first-level signal model while pipeline **6SF** applied FSL’s; in Fig. S1, pipeline **5AF** applied AFNI’s signal model while pipeline **6AF** applied FSL’s. In both cases, switching to FSL’s signal model led to a considerable change in the final thresholded results, evidenced by the sizable difference in the Dice coefficients obtained for these two specific workflows (highlighted by the blue windows in the Dice plots for Figs. 2 and S1, bottom right). Particularly, a large cluster of positive activation observed in the anterior cingulate for the pipelines implementing SPM’s and AFNI’s first-level signal model was not identified in the corresponding set of thresholded results that used FSL’s signal model (Figs. 2 and S1, thresholded maps, middle). However, these changes were not simply caused by subtle differences magnified by the thresholding, as considerable decreases in the correlation values for the unthresholded maps can also be observed for these two workflows (Figs. 2 and S1, correlation plots, bottom left), indicating that dissimilarities between the signal model’s applied by the two pipelines ultimately led to radically different activation profiles in the unthresholded group-level *t*-statistic images.

**Figure 2:**
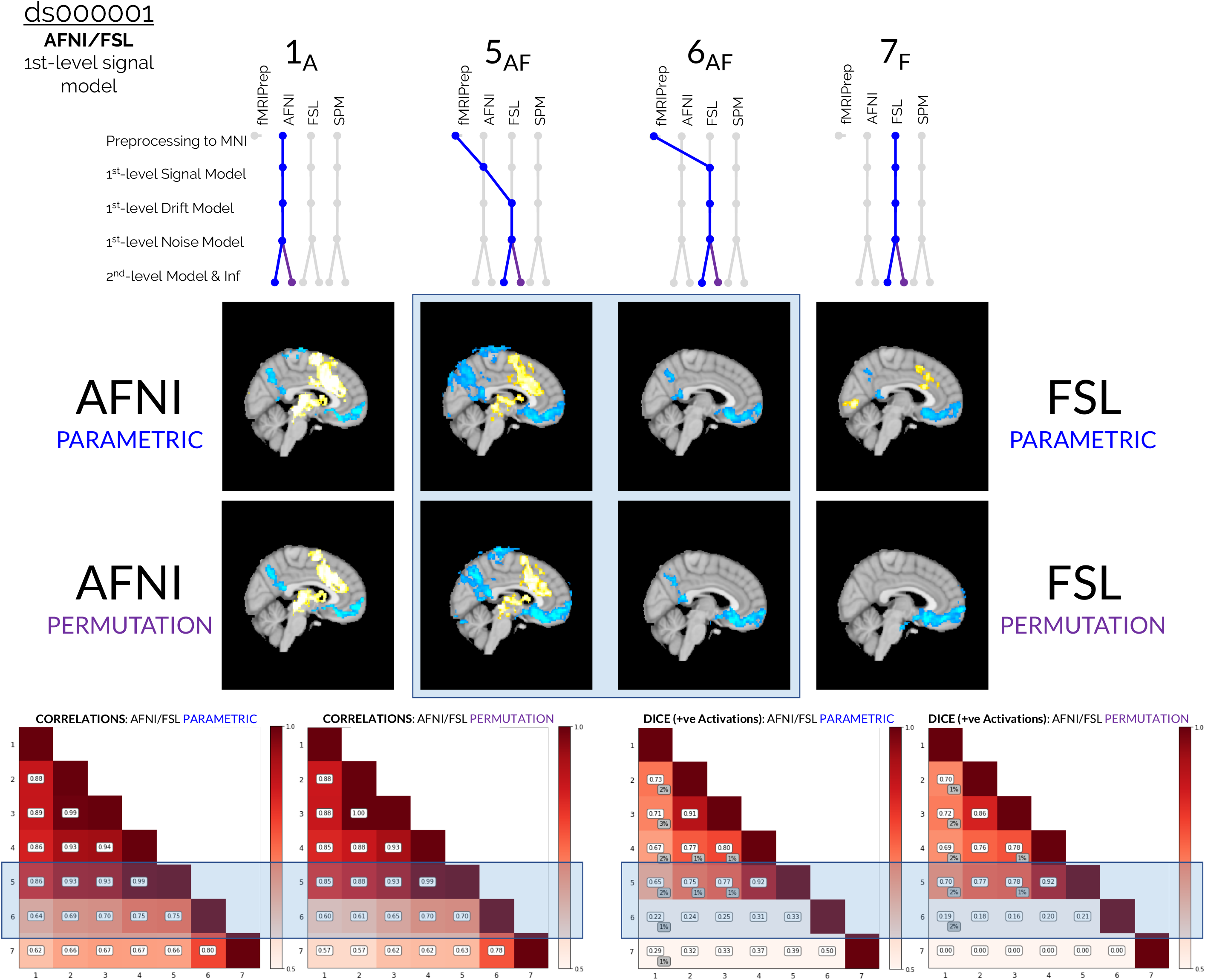
Comparisons of the group-level thresholded *t*-statistic maps, correlation values, and Dice coefficients obtained from reanalyses of the ds000001 dataset, focusing on the collection of results obtained from hybrid pipelines that implemented procedures from both AFNI and FSL. Blue windows highlight the disagreement between the two sets of results given by pipelines **5AF** and **6AF**, which differed only as to whether AFNI’s or FSL’s first-level signal model was used. The interchange of the signal model between packages led to more expansive differences in the final results than any other individual processing step: in the thresholded group-level *t*-statistic maps (middle), the expansive clusters of positive activation in the anterior cingulate (among other brain regions) identified by the pipeline using AFNI’s signal model (workflow **5AF**) were lost when interchanged with FSL’s signal model (workflow **6AF**). Differences in the thresholded maps are also reflected in the Dice coefficient matrices (bottom right), where the Dice values dramatically fell due the change of signal model between pipelines **5AF** and **6AF**. The moderate decreases also seen in the correlation values for these two pipelines (bottom-left) indicate that the interchange of signal model led to a considerable difference in the overall activation profile of the unthresholded *t*-stastistic image.

In terms of preprocessing, the results from our analyses across all studies indicate that AFNI’s preprocessing pipeline was the most similar of the three software packages to fMRIPrep, while FSL’s was the least similar. Evidence for this is seen in Figures S2 and S3, where we compare results obtained for ds000001 and ds000109, respectively, for two pairs of workflows: pipelines **1A** and **2AF**, which differ only as to whether fMRIPrep or AFNI’s preprocessing procedure was carried out, and pipelines **6AF** and **7F**, which differ only as to whether fMRIPrep or FSL’s preprocessing was applied. In each plot, differences between the results from pipelines **1A** and **2AF** have been highlighted with a blue window, while differences between **6AF** and **7F** have been highlighted with a green window. In all cases, it can be seen that the final results obtained with either fMRIPrep or AFNI’s preprocessing workflow had greater comparability than the corresponding fMRIPrep/FSL results: the final thresholded activation clusters for fMRIPrep/AFNI pipelines were more similar relative to the fMRIPrep/FSL thresholded results (Figs. S2 and S3, middle plots), and the correlation and Dice coefficients comparing pipelines **1A** and **2AF** were consistently larger than the corresponding values for pipelines **6AF** and **7F** (Figs. S2 and S3, bottom-left and bottom-right plots). The fMRIPrep/SPM Dice and correlation values can be seen in the supplementary figures (Figs. S9, S10, S13, S14 and S15); on the whole, these are slightly better than the corresponding FSL values, and slightly worse than the AFNI figures.

Aside from preprocessing, the single analysis step that caused the most variation in the ds000109 results was the first-level noise model. In Figures 3 and S4, we focus our attention on how changing from AFNI’s first-level noise model (Fig. 3) or SPM’s first-level noise model (Fig. S4) to FSL’s noise model caused a more considerable change in the final results relative to the other processing steps. In both figures, it can be seen that the correlation values (Figs. 3 and S4, bottom left plots) and Dice values (Figs. 3 and S4, bottom right plots) obtained for comparisons between pipelines **3** and **4** (which differ only by choice of first-level noise model) were generally worse than all other comparisons of pipelines varying by a single analysis step (values magnified by the blue windows in the bottom plots). However, it is notable that all correlations and Dice values were greater than 0.8 here, and the overall variation between results for ds000109 was much less than that observed for ds000001.

**Figure 3:**
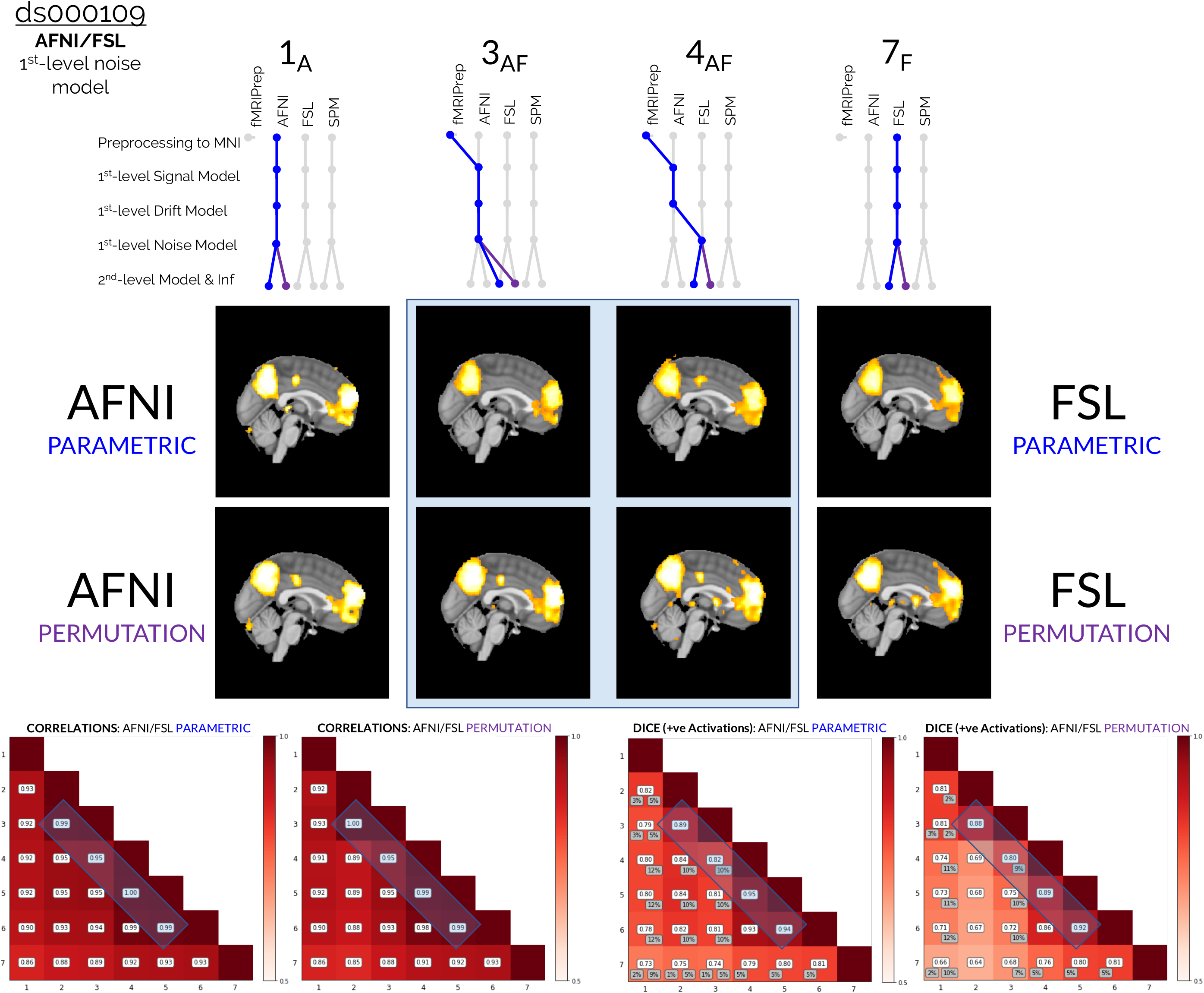
Comparisons of the group-level thresholded *t*-statistic maps, correlation values, and Dice coefficients obtained from reanalyses of the ds000109 dataset, focusing on the collection of results obtained from hybrid pipelines that implemented procedures from both AFNI and FSL. The two sets of results given by pipelines **3AF** and **4AF** are displayed, which differed only as to whether AFNI’s or FSL’s first-level noise model was implemented. Preprocessing aside, the interchange of the first-level noise model impacted the final group-level results more than any other modelling decision. This is highlighted in the correlation and Dice plots at the bottom of the figure: the blue windows on the off-diagonals show that the pairwise correlation and Dice values for pipelines **3AF** and **4AF** are smaller than the corresponding values obtained for all other pairs of adjacent pipelines. In the thresholded *t*-statistic maps (blue window, middle), it can be seen that while both of these workflows captured the main effects in the precuneus and frontal brain areas, pipeline **4AF** (that used FSL’s first-level noise model) also determined numerous smaller activation clusters which were not captured by pipeline **3AF** (that used AFNI’s noise model).

For ds000120, the group-level model and inference procedure was the largest source of variability between software. This is seen in Figure 4, where the two analysis workflows which differed only by the choice of group-level inference model are compared: pipeline **2AS** applying AFNI’s group-level modelling and inference, and pipeline **3AS** applying SPM’s group-level modelling and inference. Similar to ds000109, while the main effects were captured in the thresholded *t*-statistic maps by both packages (for ds000120, both **2AS** and **3AS** identified large clusters in the visual cortex), there was more disagreement over the presence of weaker effects. In this case, pipeline **2AS** (that used AFNI’s group-level inference model) determined a greater quantity of smaller clusters scattered across central regions of the brain compared to pipeline **3AS** (that used SPM’s group model). It is also notable that AFNI’s group-level model generally determined larger *t*-statistic values in the main activated regions compared to SPM (brighter clusters in the visual cortex for pipelines **1A** and **2AS** compared to **3AS** and **7S** in Fig. 4).

**Figure 4:**
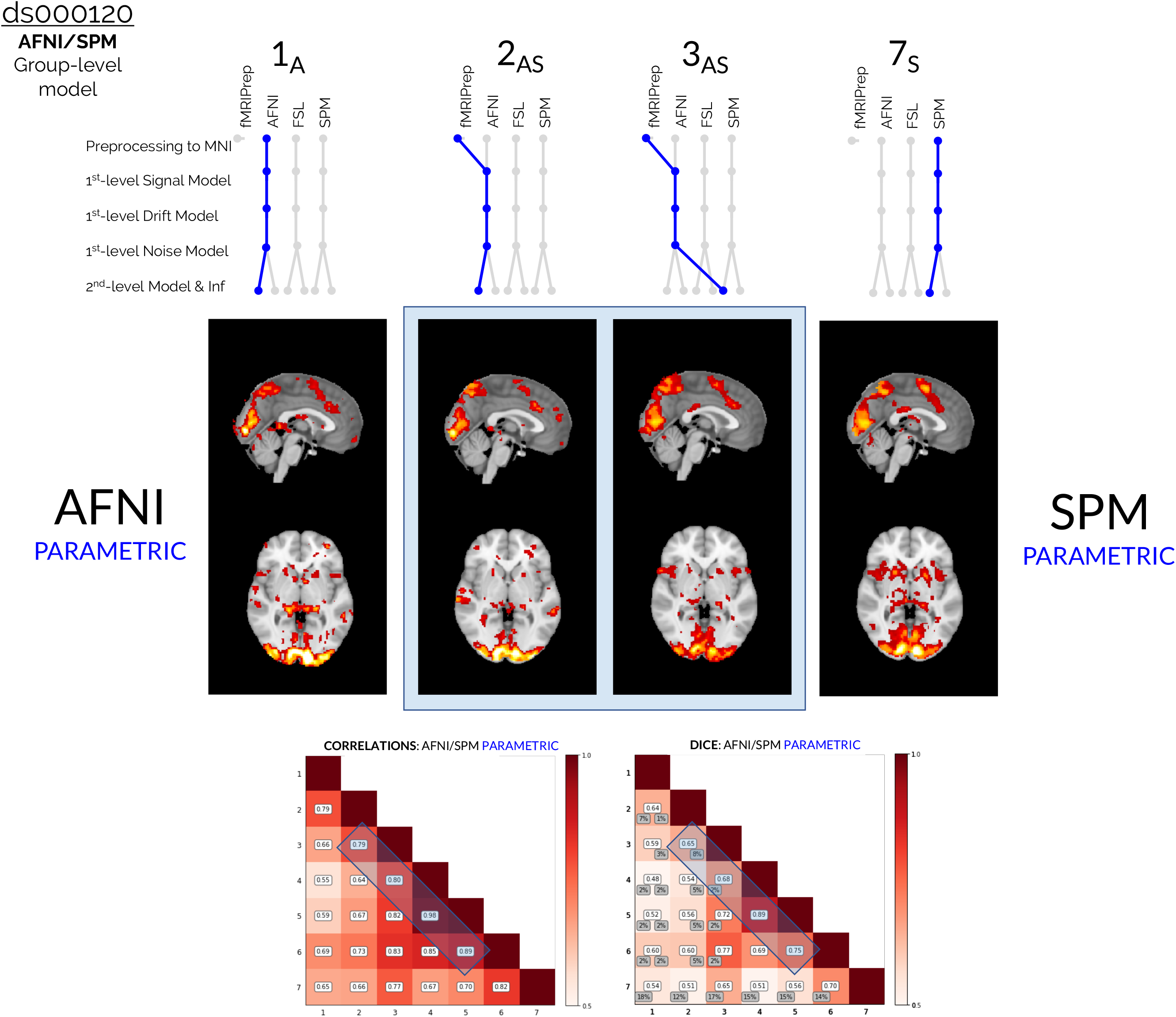
Comparisons of the group-level thresholded *t*-statistic maps, correlation values, and Dice coefficients obtained from reanalyses of the ds000120 dataset, focusing on the collection of results obtained from hybrid pipelines that implemented procedures from both AFNI and SPM. The two sets of results given by pipelines **2AS** and **3AS** are displayed, which differed only as to whether AFNI’s or SPM’s group-level inference model was implemented. The interchange of the second-level model impacted the final results more than any other modelling decision. This is highlighted in the correlation and Dice plots at the bottom of the figure: the blue windows on the off-diagonals show that the pairwise correlation and Dice values for pipelines **2AS** and **3AS** are smaller than the corresponding values obtained for all other pairs of adjacent pipelines. In the thresholded *t*-statistic maps (blue window, middle), while both pipelines captured the main effects in the visual cortex, pipeline **2AS** (that used AFNI’s group-level model and inference) identified more smaller clusters scattered across central brain regions compared to pipeline **3AS** (that used SPM’s group-level model and inference). It can also be seen that AFNI’s inference model reported larger *t*-statistic values in activated regions compared to SPM.

Finally, we observed that the choice as to which software’s first-level drift model was applied in the analysis pipeline led to minimal changes in the final analysis results. This is shown in Figures S5 and S6, where we highlight the similarity in results obtained for ds000001 and ds000109, respectively, for two workflows (pipelines **4SF** and **5SF**) which only differed as to whether SPM or FSL was used to model the drift. In both figures, it can be seen that the thresholded results obtained for these two pipelines (Figs. S5 and S6, middle plots) were qualitatively very similar, that the unthresholded maps obtained with these two workflows were almost perfectly correlated (Figs. S5 and S6, bottom-left plots), and that Dice comparisons for the thresholded maps were consistently around 90%.

### 3.2. Consensus Analyses

Slice views of the thresholded *z*-statistic maps from the consensus analyses performed on the ds000001 and ds000109 datasets are presented in Figures 5 and 6, respectively. For each dataset, the consensus analysis took the form of an image-based meta-analysis conducted on the unthresholded group-level *z*-statistic maps obtained from *all* 26 pipelines through which the data had been analyzed. The image-based meta-analysis computed a third-level *z*-statistic map, where each statistic value in the image reflected the level of evidence to which all pipelines had agreed activation was present at a given voxel. This map was then thresholded to determine voxels for which the consensus *z*-statistic was significantly greater than zero after a voxelwise FDR correction (*p* < 0.05). The equivalent one-sided correction was also performed to determine voxels whose consensus statistic was significantly *less* than zero.

**Figure 5:**
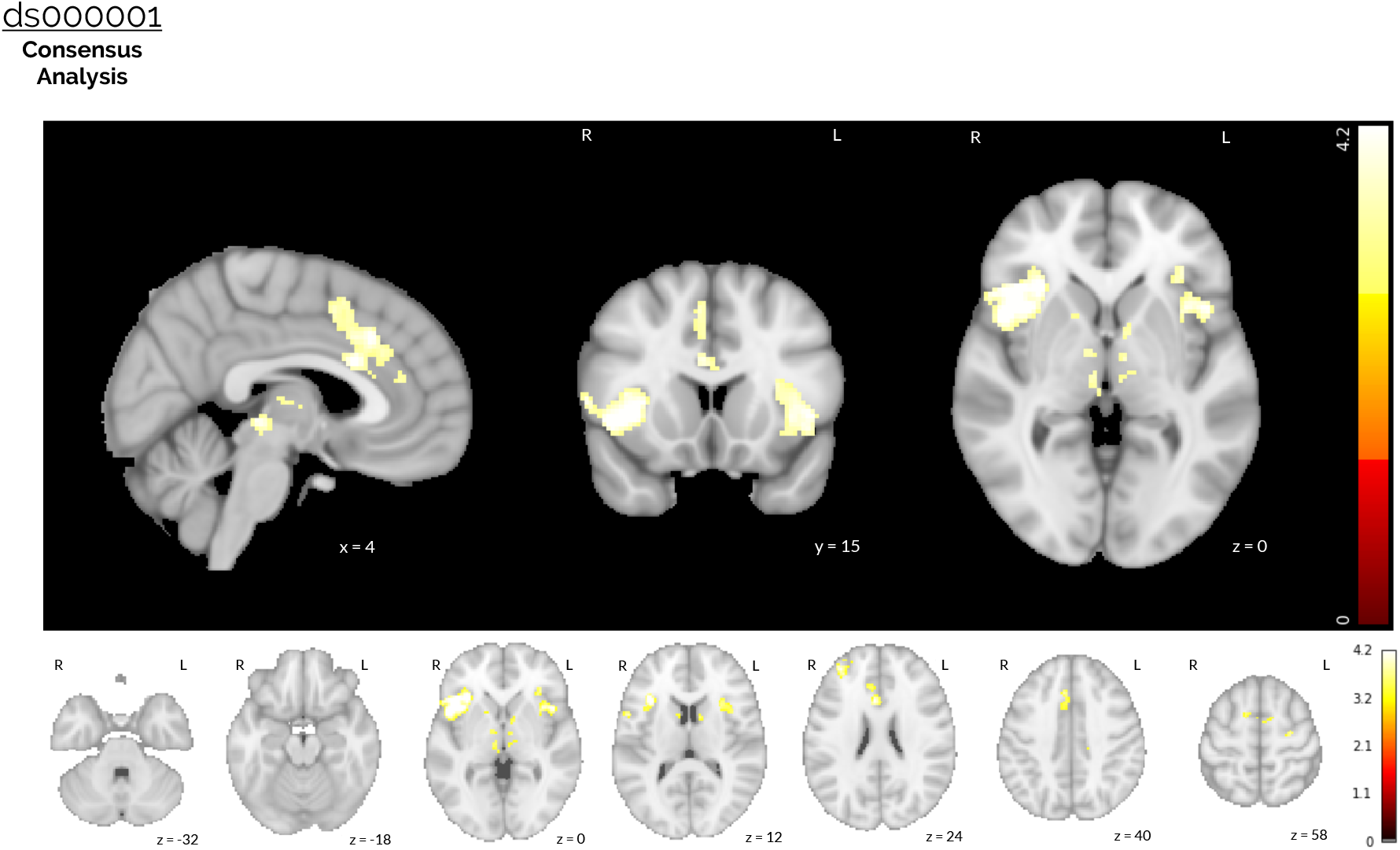
Results of the ds000001 image-based meta-analysis. A consensus analysis was performed on the unthresholded *z*-statistic maps obtained from all 26 pipelines used to analyze the ds000001 dataset, accounting for the correlation between pipelines owing to the same underlying data and identical procedures implemented across parts of the analysis workflow. The thresholded *z*-statistic map displayed shows voxels for which the group-level consensus statistic was significantly greater than zero after a voxelwise FDR correction (*p* < 0.05). Activation was found in the anterior cingulate, the insular cortex (bilateral) and the inferior frontal gyrus (right side only) after accounting for between-pipeline variation. No (negative) activation was obtained when the equivalent inference was performed to determine voxels where the consensus statistic was significantly *less* than zero.

**Figure 6:**
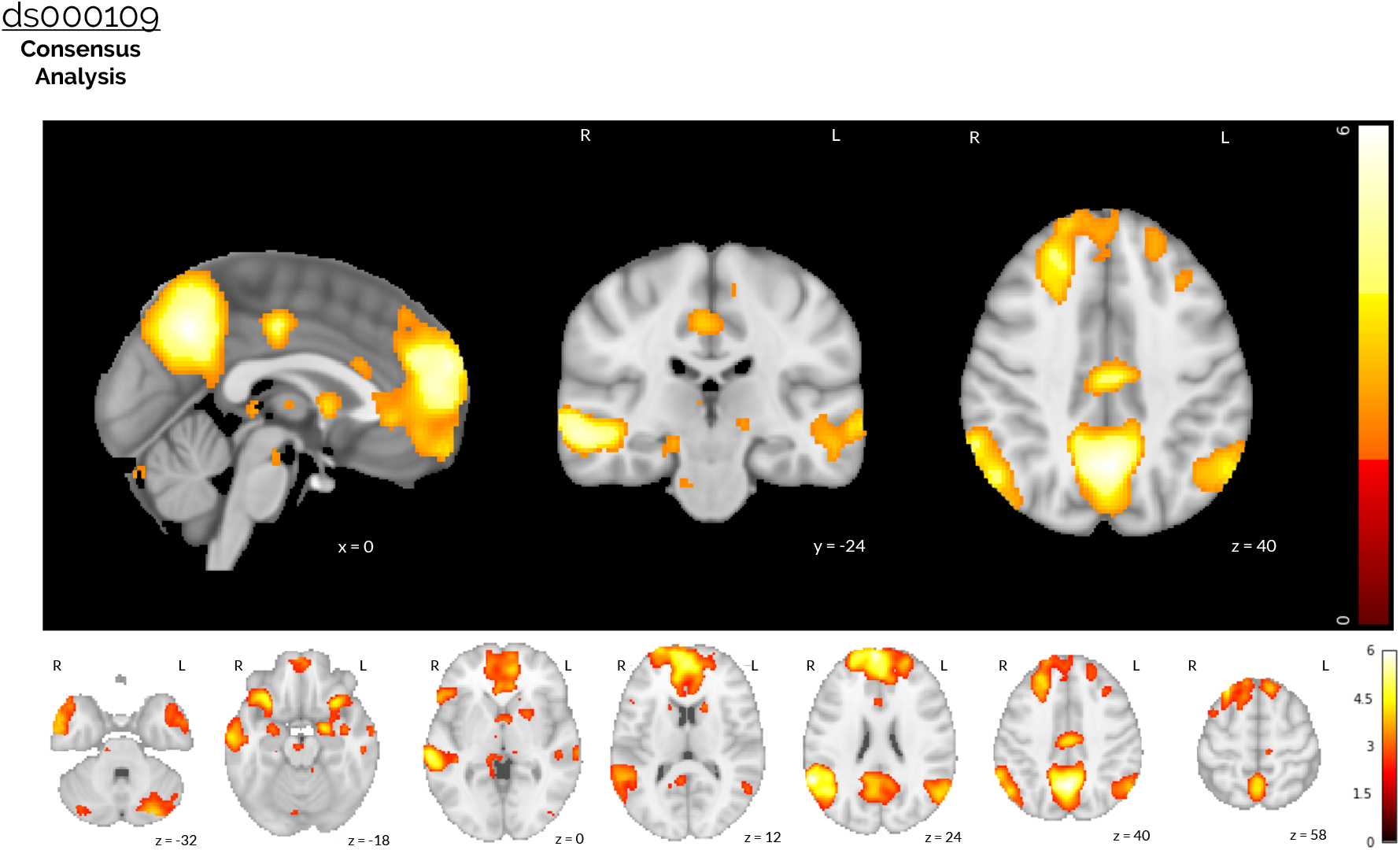
Results of the ds000109 image-based meta-analysis. A consensus analysis was performed on the unthresholded *z*-statistic statistical maps obtained from all 26 pipelines used to analyze the ds000109 dataset, accounting for the correlation between pipelines owing to the same underlying data and identical procedures implemented across parts of the analysis workflow. The thresholded *z*-statistic map displayed shows voxels for which the group-level consensus statistic was significantly greater than zero after a voxelwise FDR correction (*p* < 0.05). Large activation clusters included areas of the precuneus, frontal pole and superior frontal gyrus, the bilateral superior occipital cortex and angular gyri (bilateral). Further activation was found in the middle temporal gyrus (posterior and anterior divisions, bilateral), the left and right amygdalae, and the posterior cingulate gyrus. No (negative) activation was obtained when the equivalent inference was performed to determine voxels where the consensus statistic was significantly *less* than zero.

For ds000001, the thresholded *z*-statistic image presented in Fig. 5 shows a consensus across pipelines of positive activation in the anterior cingulate, the insular cortex (bilateral) and the inferior frontal gyrus (right side only). Significant activation was also determined in these brain areas for nearly all of the thresholded group-level *t*-statistic maps obtained from each individual analysis workflow, as can be seen in Supplementary Figs. S7-S10. However, the thresholded *z*-statistic for a consensus on negative activations failed to determine any brain areas that were statistically significant after FDR correction.

For ds000109, the thresholded *z*-statistic image presented in Fig. 6 shows a consensus across pipelines of positive activation in a variety of brain regions. Large activation clusters covered areas of the precuneus, frontal pole and superior frontal gyrus, the bilateral superior occipital cortex and angular gyri (bilateral), and further activation was determined in the middle temporal gyrus (posterior and anterior divisions, bilateral), the left and right amygdalae, and the posterior cingulate gyrus. The main effects seen here were also captured in all of the thresholded group-level *t*-statistic maps obtained from each individual analysis workflow, displayed in Supplementary Figs. S11-S14. Once again, the thresholded *z*-statistic for a consensus on negative activations failed to determine any brain areas that were statistically significant after the voxelwise correction.

## 4. Discussion

Comparisons of the statistical maps obtained from the collection of pipelines applied to the three datasets have shown both the robustness and fragility of group-level task-fMRI results in response to variation of the software package at different stages of the analysis workflow. While results were found to be highly stable across all datasets when the analysis package used to model the low-frequency fMRI drifts was interchanged, other analytic manipulations produced more appreciable changes in the group-level results. For instance, the final regions of activation obtained for the ds000001 dataset were found to be highly contingent on the software package used to model the fMRI signal; switching between AFNI/FSL’s signal model (pipelines **5AF** and **6AF** in Fig. 2) and SPM/FSL’s signal model (pipelines **5SF** and **6SF** in Fig. S1) both produced considerable differences in the *t*-statistic maps (Dice coefficients less than 0.35 for comparisons of the thresholded maps, correlations less than 0.75 for unthresholded maps). However, for ds000109 the change of signal model had minimal impact on the final results (in Supplementary Fig. S11, comparisons of the **5AF** and **6AF** unthresholded maps yielded a correlation of 0.99, Dice coefficient of 0.94 for positive activations). In contrast, the interchange of ds000109’s first-level noise model produced greater relative differences, and for ds000120, the group-level model was found to be the largest modelling source of between-software variability (Figs. 3 and S4, Supplementary Figs. S11-S15).

Importantly, these results do not provide an indication as to which software package is better or worse; without a gold standard to compare to, no such claims can be made. However, these findings do suggest that the main sources of software-variability across the analysis pipeline can be heterogeneous and dependent on external factors such as the analysis design or task paradigm under investigation. One reason that the quantitative comparisons for the ds000001 dataset were generally worse than the corresponding ds000109 and ds000120 comparisons is likely to be due to the smaller sample size for this study (15 for ds000001 vs 21 and 17 for ds000109 and ds000120 respectively). As well as this, the larger impact of the signal model for ds000001 may be attributed to varying aspects between the three studies’ analysis designs. In particular, the event-related design used for ds000001’s balloon analog risk task could have been more sensitive to differences between each package’s hemodyanmic response model compared to the block design used for ds000109. In addition to this, while ds000109 and ds000120 did not apply any modulated regressor orthogonalization methods, for each of the three ds000001 task events represented in the GLM (pumps, cash-outs, and explosions) the response time regressors were orthogonalized with respect to the average activity regressor (e.g. *pumps_response_time_* condition orthogonalized with respect to *pumps_average_* condition). It has been previously observed in the fMRI literature that the three software packages handle orthogonalization differently (Mumford et al., 2015): while in FSL each regressor can be orthogonalized with respect to any other individual regressor the user has specified, in SPM orthogonalization is applied automatically, and each regressor is orthogonalized with respect to *all* other conditions preceding it in the model. We suspect that differences in the shared variance between regressors caused by divergent orthoganlization procedures across the three packages is one of the reasons that the choice of signal model proved to be so influential for ds0000001.

The inference procedure carried out specifically for ds000001 may have also contributed to variation in the activation clusters identified in the thresholded *t*-statistic maps, particularly for this studies’ collection of parametric inference results. Group-level inference was conducted using a cluster-forming threshold of *p* < 0.01 uncorrected for the ds000001 study. However, in Eklund et al. (2016) it was found that parametric clusterwise inference using this cluster-forming threshold led to varying degrees of inflated false activations across the three software packages. Notably, while these findings may in-part explain the poor Dice coefficients for the ds000001 parametric results, they do not have any bearing on the correlation comparisons (since correlations were comparisons of the *unthresholded* maps) or the collection of corresponding nonparametric results (since permutation inference was shown to perform as expected in Eklund et al.). These findings also do not affect the ds000001 consensus analysis results in Section 3.2, which used a voxelwise FDR correction for inference. Since the correlation and nonparametric inference comparisons were observed to be poorer for ds000001 compared to the other studies too, this could suggest that divergence between the parametric results were also caused by other factors than the cluster-forming threshold.

From the quantitative comparisons presented for all studies in Figs. 2-S6, it is notable how seemingly small differences in the unthresholded maps could be amplified after thresholding. Even when pairwise correlations of the unthresholded statistic maps were considerably high, in many cases the corresponding Dice comparisons measuring the overlap of activation in the thresholded maps were substantially lower. This is illustrated by the comparisons of pipelines **6AF** and **7F** in Fig. S3 (and Supplementary Fig. S11), where the correlation between these two pipelines’ unthresholded *t*-statistic maps was 0.93, but the Dice coefficient for negative activations in the thresholded maps was 0. Ultimately, this was due to pipeline **6AF** identifying two clusters of negative activation in the left and right interior temporal gyri, while pipeline **7F** didn’t determine any negative activation. The overriding issue here is the dichotomous nature of thresholding; because maps are binarized into regions of activation and non-activation based on a single cut-off value, substantially different thresholded maps can be obtained depending on whether a cluster’s size is marginally above or below the threshold. This is a particularly salient reason why we believe the unthresholded statistical maps should always be shared. Access to the unthresholded maps enables further meta-analyses of the data to be conducted, where the variation of clusters across diverse samples (and analysis workflows) can be quantified in order to determine where results converge. The consensus analyses carried out as part of this work exemplify the benefits to such an approach, and notably the thresholded consensus map for ds000109 (Fig. 6) didn’t identify any regions of negative activation after accounting for the inter-pipeline variation between individual results.

In conclusion, we believe that multi-software analyses are essential to understanding the nature and origins of inter-software differences. For pipeline elements that produce the greatest differences, further study will be required to determine an optimal or preferred method (Churchill et al., 2015). However, until further research on pipeline harmonisation has been carried out, the field will depend on traditional replication analyses. To this end, it is vital that both practitioners and publishers embrace the importance of replication studies and the publication of null findings. Alongside this, replication can only become possible if data sharing practices become commonplace in the field. In this work, we have shared all of our statistical results (both unthreshold and thresholded maps) and analysis code via public online repositories (Neurovault and Github/Zenodo), and we hope that other researchers will follow suit to advance transparency in neuroimaging science.

## 5. Data Sharing

All scripts and results have been made available through our Open Science Framework (OSF; Foster and Deardorff, 2017) Project at https://osf.io/axy3w/ (Bowring et al., 2021), and all group-level statistic maps have been made available on NeuroVault: https://neurovault.org/collections/8381/, https://neurovault.org/collections/7113/, https://neurovault.org/collections/9324/, for ds0000001, ds000109, and ds000120, respectively. Python Jupyter Notebooks have also been shared for each of the three studies, harvesting the results data from Neurovault and applying the comparison methods discussed in the previous section to create all the figures used here. All analysis scripts, results reports, and notebooks for each study are available through Zenodo (Nielsen et al., 2014) at http://doi.org/10.5281/zenodo.5070414.

## 6. Acknowledgements

All authors would like to thank Adam Huffman for his monumental efforts to install the software required for this work in the midst of a pandemic. We would also like to thank the fMRIPrep and NeuroStars communities, and particularly, Oscar Esteban, Franklin Feingold, Ross Blair, and Russ Poldrack, for their assistance with fMRIPrep. This study would not have been possible without OpenNeuro and the pool of researchers who have contributed their data to the project, to whom we are grateful.

## Notes

https://osf.io/axy3w/

https://neurovault.org/collections/8381/

https://neurovault.org/collections/7113/

https://neurovault.org/collections/9324/

http://doi.org/10.5281/zenodo.5070414

### Competing Interest Statement

The authors have declared no competing interest.

## References

Brian B Avants, Charles L Epstein, Murray Grossman, and James C Gee. Symmetric diffeomorphic image registration with cross-correlation: evaluating automated labeling of elderly and neurodegenerative brain. Medical image analysis, 12(1):26–41, 2008.

Yashar Behzadi, Khaled Restom, Joy Liau, and Thomas T Liu. A component based noise correction method (compcor) for bold and perfusion based fmri. Neuroimage, 37(1):90–101, 2007.

Yoav Benjamini, Abba M Krieger, and Daniel Yekutieli. Adaptive linear step-up procedures that control the false discovery rate. Biometrika, 93(3):491–507, 2006.

Rotem Botvinik-Nezer, Felix Holzmeister, Colin F Camerer, Anna Dreber, Juergen Huber, Magnus Johannesson, Michael Kirchler, Roni Iwanir, Jeanette A Mumford, R Alison Adcock, et al. Variability in the analysis of a single neuroimaging dataset by many teams. Nature, pages 1–7, 2020.

Alex Bowring, Thomas Nichols, and Camille Maumet. Identifying sources of software-dependent differences in task fmri analyses, Jun 2021. URL osf.io/axy3w.

Alexander Bowring, Camille Maumet, and Thomas E. Nichols. Exploring the impact of analysis software on task fmri results. Human Brain Mapping, 40(11):3362–3384, 2019. doi: 10.1002/hbm.24603. URL https://onlinelibrary.wiley.com/doi/abs/10.1002/hbm.24603.

Dana Carney. My position on “power poses”, 2017. URL http://faculty.haas.berkeley.edu/dana_carney/pdf_my%20position%20on%20power%20poses.pdf. Accessed: 2020-11-13.

Dana R. Carney, Amy J.C. Cuddy, and Andy J. Yap. Power posing: Brief nonverbal displays affect neuroendocrine levels and risk tolerance. Psychological Science, 21(10):1363–1368, 2010. doi: 10.1177/0956797610383437. URL https://doi.org/10.1177/0956797610383437. PMID: 20855902.

Joshua Carp. On the plurality of (methodological) worlds: Estimating the analytic flexibility of fmri experiments. Frontiers in Neuroscience, 6:149, 2012. ISSN 1662-453X. doi: 10.3389/fnins.2012.00149. URL https://www.frontiersin.org/article/10.3389/fnins.2012.00149.

Nathan W. Churchill, Robyn Spring, Babak Afshin-Pour, Fan Dong, and Stephen C. Strother. An automated, adaptive framework for optimizing preprocessing pipelines in task-based functional mri. PLOS ONE, 10(7): 1–25, 07 2015. doi: 10.1371/journal.pone.0131520. URL https://doi.org/10.1371/journal.pone.0131520.

Open Science Collaboration. Estimating the reproducibility of psychological science. Science, 349(6251), 2015. ISSN 0036-8075. doi: 10.1126/science.aac4716. URL https://science.sciencemag.org/content/349/6251/aac4716.

Anders M Dale, Bruce Fischl, and Martin I Sereno. Cortical surface-based analysis: I. segmentation and surface reconstruction. Neuroimage, 9(2):179–194, 1999.

Charles R Ebersole, Olivia E Atherton, Aimee L Belanger, Hayley M Skulborstad, Jill M Allen, Jonathan B Banks, Erica Baranski, Michael J Bernstein, Diane B V Bonfiglio, Leanne Boucher, Elizabeth R Brown, Nancy I Budiman, Athena H Cairo, Colin A Capaldi, Christopher R Chartier, Joanne M Chung, David C Cicero, Jennifer A Coleman, John G Conway, William E Davis, Thierry Devos, Melody M Fletcher, Komi German, Jon E Grahe, Anthony D Hermann, Joshua A Hicks, Nathan Honeycutt, Brandon Humphrey, Matthew Janus, David J Johnson, Jennifer A Joy-Gaba, Hannah Juzeler, Ashley Keres, Diana Kinney, Jacqeline Kirshenbaum, Richard A Klein, Richard E Lucas, Christopher J N Lustgraaf, Daniel Martin, Madhavi Menon, Mitchell Metzger, Jaclyn M Moloney, Patrick J Morse, Radmila Prislin, Timothy Razza, Daniel E Re, Nicholas O Rule, Donald F Sacco, Kyle Sauerberger, Emily Shrider, Megan Shultz, Courtney Siemsen, Karin Sobocko, R Weylin Sternglanz, Amy Summerville, Konstantin O Tskhay, Zack van Allen, Leigh Ann Vaughn, Ryan J Walker, Ashley Weinberg, John Paul Wilson, James H Wirth, Jessica Wortman, and Brian A Nosek. Many labs 3: Evaluating participant pool quality across the academic semester via replication. J. Exp. Soc. Psychol., 67: 68–82, November 2016.

Anders Eklund, Thomas E Nichols, and Hans Knutsson. Cluster failure: Why fmri inferences for spatial extent have inflated false-positive rates. Proceedings of the national academy of sciences, 113(28):7900–7905, 2016.

Oscar Esteban, Christopher J Markiewicz, Ross W Blair, Craig A Moodie, A Ilkay Isik, Asier Erramuzpe, James D Kent, Mathias Goncalves, Elizabeth DuPre, Madeleine Snyder, et al. fmriprep: a robust preprocessing pipeline for functional mri. Nature methods, 16(1):111–116, 2019.

Oscar Esteban, Christopher J. Markiewicz, Christopher Burns, Mathias Goncalves, Dorota Jarecka, Erik Ziegler, Shoshana Berleant, David Gage Ellis, Basile Pinsard, Cindee Madison, Michael Waskom, Michael Philipp Notter, Daniel Clark, Alexandre Manhães-Savio, Dav Clark, Kesshi Jordan, Michael Dayan, Yaroslav O. Halchenko, Fred Loney, Taylor Salo, Blake E Dewey, Hans Johnson, Salma Bougacha, Anisha Keshavan, Benjamin Yvernault, Carlo Hamalainen, Horea Christian, Rastko Ćirić, Mathieu Dubois, Michael Joseph, Ben Cipollini, Steven Tilley II, Matteo Visconti di Oleggio Castello, Jason Wong, Alejandro De La Vega, Jakub Kaczmarzyk, Julia M. Huntenburg, Michael G. Clark, Erin Benderoff, Drew Erickson, James D. Kent, Michael Hanke, Steven Giavasis, Brendan Moloney, B. Nolan Nichols, Rosalia Tungaraza, Caroline Frohlich, Demian Wassermann, Gilles de Hollander, Arman Eshaghi, Jarrod Millman, Matteo Mancini, Dylan M. Nielson, Gael Varoquaux, Aimi Watanabe, David Mordom, Jérémy Guillon, Serge Koudoro, Andrey Chetverikov, Ariel Rokem, Benjamin Acland, Jessica Forbes, Ross Markello, Ashley Gillman, Xiang-Zhen Kong, Daniel Geisler, John Salvatore, Alexandre Gramfort, Anna Doll, Colin Buchanan, Elizabeth DuPre, Siqi Liu, Alexander Schaefer, Jens Kleesiek, Sharad Sikka, Yannick Schwartz, John A. Lee, Aaron Mattfeld, Adam Richie-Halford, Franz Liem, Martin Felipe Perez-Guevara, Anibal Sólon Heinsfeld, Christian Haselgrove, Joke Durnez, Leonie Lampe, Russell Poldrack, Tristan Glatard, Alejandro Tabas, Chad Cumba, Fernando Pérez-García, Ross Blair, Shariq Iqbal, David Welch, William Triplett, Ali Ghayoor, R. Cameron Craddock, Carlos Correa, Dimitri Papadopoulos Orfanos, Jörg Stadler, Joshua Warner, Lucinda M. Sisk, Marcel Falkiewicz, Paul Sharp, Simon Rothmei, Sin Kim, Alejandro Weinstein, Ari E. Kahn, Erik Kastman, Katherine Bottenhorn, Martin Grignard, L. Nathan Perkins, Oliver Contier, Dale Zhou, Dmytro Bielievtsov, Gavin Cooper, Hrvoje Stojic, Janosch Linkersdörfer, Lea Waller, Mandy Renfro, Oliver Hinds, Olivia Stanley, René Küttner, Wolfgang M. Pauli, Daniel Glen, Adam Kimbler, Benjamin Meyers, Claire Tarbert, Daniel Ginsburg, Daniel Haehn, Daniel S. Margulies, Feilong Ma, Ian B. Malone, Lukas Snoek, Matthew Brett, Matthew Cieslak, Michael Hallquist, Miguel Molina-Romero, Murat Bilgel, Nat Lee, Souheil Inati, Stephan Gerhard, Sulantha Mathotaarachchi, Victor Saase, Andrew Van, Christopher John Steele, Eduard Ort, Eric Condamine, Garikoitz Lerma-Usabiaga, Isaac Schwabacher, Jaime Arias, Jeff Lai, John Pellman, Jordi Huguet, Junhao Wen, Katrin Leinweber, Kshitij Chawla, Leon Weninger, Marc Modat, Robbert Harms, Sami Kristian Andberg, Zvi Baratz, K Matsubara, Abel A. González Orozco, Ana Marina, Andrew Davison, Andrew Floren, Anne Park, Brian Cheung, Conor McDermottroe, Daniel McNamee, Dmitry Shachnev, Guillaume Flandin, Ivan Gonzalez, Jan Varada, Kai Schlamp, Kornelius Podranski, Lijie Huang, Maxime Noel, Michael R. Crusoe, Nicolas Pannetier, Ranjeet Khanuja, Sebastian Urchs, Thomas Nickson, Lijie Huang, William Broderick, Arielle Tambini, Paul Glad Mihai, Krzysztof J. Gorgolewski, and Satrajit Ghosh. nipy/nipype: 1.5.1, September 2020a. URL https://doi.org/10.5281/zenodo.4035081.

Oscar Esteban, Christopher J. Markiewicz, Mathias Goncalves, Elizabeth DuPre, James D. Kent, Taylor Salo, Rastko Ciric, Basile Pinsard, Ross W. Blair, Russell A. Poldrack, and Krzysztof J. Gorgolewski. fMRIPrep: a robust preprocessing pipeline for functional MRI, November 2020b. URL https://doi.org/10.5281/zenodo.4252786.

Vladimir S Fonov, Alan C Evans, Robert C McKinstry, CR Almli, and DL Collins. Unbiased nonlinear average age-appropriate brain templates from birth to adulthood. NeuroImage, (47):S102, 2009.

Erin D Foster and Ariel Deardorff. Open science framework (osf). Journal of the Medical Library Association: JMLA, 105(2):203, 2017.

Katie E. Garrison, David Tang, and Brandon J. Schmeichel. Embodying power: A preregistered replication and extension of the power pose effect. Social Psychological and Personality Science, 7(7):623–630, 2016. doi: 10.1177/1948550616652209. URL https://doi.org/10.1177/1948550616652209.

Krzysztof Gorgolewski, Christopher D Burns, Cindee Madison, Dav Clark, Yaroslav O Halchenko, Michael L Waskom, and Satrajit S Ghosh. Nipype: a flexible, lightweight and extensible neuroimaging data processing framework in python. Frontiers in neuroinformatics, 5:13, 2011.

Krzysztof Gorgolewski, Oscar Esteban, Gunnar Schaefer, Brian Wandell, and Russell Poldrack. Openneuro—a free online platform for sharing and analysis of neuroimaging data. Organization for human brain mapping. Vancouver, Canada, 1677(2), 2017.

Krzysztof J Gorgolewski, Tibor Auer, Vince D Calhoun, R Cameron Craddock, Samir Das, Eugene P Duff, Guillaume Flandin, Satrajit S Ghosh, Tristan Glatard, Yaroslav O Halchenko, Daniel A Handwerker, Michael Hanke, David Keator, Xiangrui Li, Zachary Michael, Camille Maumet, B Nolan Nichols, Thomas E Nichols, John Pellman, Jean-Baptiste Poline, Ariel Rokem, Gunnar Schaefer, Vanessa Sochat, William Triplett, Jessica A Turner, Gaël Varoquaux, and Russell A Poldrack. The brain imaging data structure, a format for organizing and describing outputs of neuroimaging experiments. Sci Data, 3:160044, June 2016.

Douglas N Greve and Bruce Fischl. Accurate and robust brain image alignment using boundary-based registration. Neuroimage, 48(1):63–72, 2009.

John P. A. Ioannidis. Why most published research findings are false. PLOS Medicine, 2(8):null, 08 2005. doi: 10.1371/journal.pmed.0020124. URL https://doi.org/10.1371/journal.pmed.0020124.

Mark Jenkinson, Peter Bannister, Michael Brady, and Stephen Smith. Improved optimization for the robust and accurate linear registration and motion correction of brain images. Neuroimage, 17(2):825–841, 2002.

Arno Klein, Satrajit S Ghosh, Forrest S Bao, Joachim Giard, Yrjö Häme, Eliezer Stavsky, Noah Lee, Brian Rossa, Martin Reuter, Elias Chaibub Neto, et al. Mindboggling morphometry of human brains. PLoS computational biology, 13(2):e1005350, 2017.

Richard Klein, Kate Ratliff, Michelangelo Vianello, Reginald Adams, Jr, Stěpán Bahník, Michael Bernstein, Konrad Bocian, Mark Brandt, Beach Brooks, Claudia Brumbaugh, and Others. Data from investigating variation in replicability: A “many labs” replication project. Journal of Open Psychology Data, 2(1), 2014.

Richard A Klein, Michelangelo Vianello, Fred Hasselman, Byron G Adams, Reginald B Adams, Sinan Alper, Mark Aveyard, Jordan R Axt, Mayowa T Babalola, Štěpán Bahník, Rishtee Batra, Mihály Berkics, Michael J Bernstein, Daniel R Berry, Olga Bialobrzeska, Evans Dami Binan, Konrad Bocian, Mark J Brandt, Robert Busching, Anna Cabak Rédei, Huajian Cai, Fanny Cambier, Katarzyna Cantarero, Cheryl L Carmichael, Francisco Ceric, Jesse Chandler, Jen-Ho Chang, Armand Chatard, Eva E Chen, Winnee Cheong, David C Cicero, Sharon Coen, Jennifer A Coleman, Brian Collisson, Morgan A Conway, Katherine S Corker, Paul G Curran, Fiery Cushman, Zubairu K Dagona, Ilker Dalgar, Anna Dalla Rosa, William E Davis, Maaike de Bruijn, Leander De Schutter, Thierry Devos, Marieke de Vries, Canay Doǧulu, Nerisa Dozo, Kristin Nicole Dukes, Yarrow Dunham, Kevin Durrheim, Charles R Ebersole, John E Edlund, Anja Eller, Alexander Scott English, Carolyn Finck, Natalia Frankowska, Miguel-Ángel Freyre, Mike Friedman, Elisa Maria Galliani, Joshua C Gandi, Tanuka Ghoshal, Steffen R Giessner, Tripat Gill, Timo Gnambs, Ángel Gómez, Roberto González, Jesse Graham, Jon E Grahe, Ivan Grahek, Eva G T Green, Kakul Hai, Matthew Haigh, Elizabeth L Haines, Michael P Hall, Marie E Heffernan, Joshua A Hicks, Petr Houdek, Jeffrey R Huntsinger, Ho Phi Huynh, Hans IJzerman, Yoel Inbar, Åse H Innes-Ker, William Jiménez-Leal, Melissa-Sue John, Jennifer A Joy-Gaba, Roza G Kamiloǧlu, Heather Barry Kappes, Serdar Karabati, Haruna Karick, Victor N Keller, Anna Kende, Nicolas Kervyn, Goran Knežević, Carrie Kovacs, Lacy E Krueger, German Kurapov, Jamie Kurtz, Daniël Lakens, Ljiljana B Lazarević, Carmel A Levitan, Neil A Lewis, Samuel Lins, Nikolette P Lipsey, Joy E Losee, Esther Maassen, Angela T Maitner, Winfrida Malingumu, Robyn K Mallett, Satia A Marotta, Janko Mejedović, Fernando Mena-Pacheco, Taciano L Milfont, Wendy L Morris, Sean C Murphy, Andriy Myachykov, Nick Neave, Koen Neijenhuijs, Anthony J Nelson, Félix Neto, Austin Lee Nichols, Aaron Ocampo, Susan L O’Donnell, Haruka Oikawa, Masanori Oikawa, Elsie Ong, Gábor Orosz, Malgorzata Osowiecka, Grant Packard, Rolando Pérez-Sánchez, Boban Petrović, Ronaldo Pilati, Brad Pinter, Lysandra Podesta, Gabrielle Pogge, Monique M H Pollmann, Abraham M Rutchick, Patricio Saavedra, Alexander K Saeri, Erika Salomon, Kathleen Schmidt, Felix D Schönbrodt, Maciej B Sekerdej, David Sirlopú, Jeanine L M Skorinko, Michael A Smith, Vanessa Smith-Castro, Karin C H J Smolders, Agata Sobkow, Walter Sowden, Philipp Spachtholz, Manini Srivastava, Troy G Steiner, Jeroen Stouten, Chris N H Street, Oskar K Sundfelt, Stephanie Szeto, Ewa Szumowska, Andrew C W Tang, Norbert Tanzer, Morgan J Tear, Jordan Theriault, Manuela Thomae, David Torres, Jakub Traczyk, Joshua M Tybur, Adrienn Ujhelyi, Robbie C M van Aert, Marcel A L M van Assen, Marije van der Hulst, Paul A M van Lange, Anna Elisabeth van’t Veer, Alejandro Vásquez-Echeverría, Leigh Ann Vaughn, Alexandra Vázquez, Luis Diego Vega, Catherine Verniers, Mark Verschoor, Ingrid P J Voermans, Marek A Vranka, Cheryl Welch, Aaron L Wichman, Lisa A Williams, Michael Wood, Julie A Woodzicka, Marta K Wronska, Liane Young, John M Zelenski, Zeng Zhijia, and Brian A Nosek. Many labs 2: Investigating variation in replicability across samples and settings. Advances in Methods and Practices in Psychological Science, 1(4):443–490, December 2018.

Cornelius Lanczos. Evaluation of noisy data. Journal of the Society for Industrial and Applied Mathematics, Series B: Numerical Analysis, 1(1):76–85, 1964.

Scott E Maxwell, Michael Y Lau, and George S Howard. Is psychology suffering from a replication crisis? what does “failure to replicate” really mean? Am. Psychol., 70(6):487–498, September 2015.

Joseph M Moran, Eshin Jolly, and Jason P Mitchell. Social-cognitive deficits in normal aging. J. Neurosci., 32 (16):5553–5561, April 2012.

Jeanette A Mumford, Jean-Baptiste Poline, and Russell A Poldrack. Orthogonalization of regressors in fmri models. PloS one, 10(4):e0126255, 2015.

Lars Holm Nielsen, Tim Smith, Chris Erdmann, and Tibor Simko. Zenodo-an open dependable home for the long-tail of science. Open Repositories 2014, 2014.

Aarthi Padmanabhan, Charles F Geier, Sarah J Ordaz, Theresa Teslovich, and Beatriz Luna. Developmental changes in brain function underlying the influence of reward processing on inhibitory control. Dev. Cogn. Neurosci., 1(4):517–529, October 2011.

Jonathan D Power, Anish Mitra, Timothy O Laumann, Abraham Z Snyder, Bradley L Schlaggar, and Steven E Petersen. Methods to detect, characterize, and remove motion artifact in resting state fmri. Neuroimage, 84: 320–341, 2014.

Eva Ranehill, Anna Dreber, Magnus Johannesson, Susanne Leiberg, Sunhae Sul, and Roberto A. Weber. Assessing the robustness of power posing: No effect on hormones and risk tolerance in a large sample of men and women. Psychological Science, 26(5):653–656, 2015. doi: 10.1177/0956797614553946. URL https://doi.org/10.1177/0956797614553946. PMID: 25810452.

Theodore D Satterthwaite, Mark A Elliott, Raphael T Gerraty, Kosha Ruparel, James Loughead, Monica E Calkins, Simon B Eickhoff, Hakon Hakonarson, Ruben C Gur, Raquel E Gur, et al. An improved framework for confound regression and filtering for control of motion artifact in the preprocessing of resting-state functional connectivity data. Neuroimage, 64:240–256, 2013.

Tom Schonberg, Craig R Fox, Jeanette A Mumford, Eliza Congdon, Christopher Trepel, and Russell A Poldrack. Decreasing ventromedial prefrontal cortex activity during sequential risk-taking: an FMRI investigation of the balloon analog risk task. Front. Neurosci., 6:80, June 2012.

Joseph P. Simmons, Leif D. Nelson, and Uri Simonsohn. False-positive psychology: Undisclosed flexibility in data collection and analysis allows presenting anything as significant. Psychological Science, 22(11):1359–1366, 2011. doi: 10.1177/0956797611417632. URL https://doi.org/10.1177/0956797611417632. PMID: 22006061.

Kristopher M Smith and Coren L Apicella. Winners, losers, and posers: The effect of power poses on testosterone and risk-taking following competition. Hormones and Behavior, 92:172 – 181, 2017. ISSN 0018-506X. doi: https://doi.org/10.1016/j.yhbeh.2016.11.003. URL http://www.sciencedirect.com/science/article/pii/S0018506X1630023X. Hormones and Human Competition.

Fritz Strack, Leonard Martin, and Sabine Stepper. Inhibiting and facilitating conditions of the human smile: A nonobtrusive test of the facial feedback hypothesis. Journal of personality and social psychology, 54:768–77, 06 1988. doi: 10.1037/0022-3514.54.5.768.

Denes Szucs and John P. A. Ioannidis. Empirical assessment of published effect sizes and power in the recent cognitive neuroscience and psychology literature. PLOS Biology, 15(3):1–18, 03 2017. doi: 10.1371/journal.pbio.2000797. URL https://doi.org/10.1371/journal.pbio.2000797.

Nicholas J Tustison, Brian B Avants, Philip A Cook, Yuanjie Zheng, Alexander Egan, Paul A Yushkevich, and James C Gee. N4itk: improved n3 bias correction. IEEE transactions on medical imaging, 29(6):1310–1320, 2010.

E.-J. Wagenmakers, T. Beek, L. Dijkhoff, Q. F. Gronau, A. Acosta, Jr. R. B. Adams, D. N. Albohn, E. S. Allard, S. D. Benning, E.-M. Blouin-Hudon, L. C. Bulnes, T. L. Caldwell, R. J. Calin-Jageman, C. A. Capaldi, N. S. Carfagno, K. T. Chasten, A. Cleeremans, L. Connell, J. M. DeCicco, K. Dijkstra, A. H. Fischer, F. Foroni, U. Hess, K. J. Holmes, J. L. H. Jones, O. Klein, C. Koch, S. Korb, P. Lewinski, J. D. Liao, S. Lund, J. Lupianez, D. Lynott, C. N. Nance, S. Oosterwijk, A. A. Ozdoǧru, A. P. Pacheco-Unguetti, B. Pearson, C. Powis, S. Riding, T.-A. Roberts, R. I. Rumiati, M. Senden, N. B. Shea-Shumsky, K. Sobocko, J. A. Soto, T. G. Steiner, J. M. Talarico, Z. M. van Allen, M. Vandekerckhove, B. Wainwright, J. F. Wayand, R. Zeelenberg, E. E. Zetzer, and R. A. Zwaan. Registered replication report: Strack, martin, & stepper (1988). Perspectives on Psychological Science, 11(6):917–928, 2016. doi: 10.1177/1745691616674458. URL https://doi.org/10.1177/1745691616674458. PMID: 27784749.

Yongyue Zhang, Michael Brady, and Stephen Smith. Segmentation of brain mr images through a hidden markov random field model and the expectation-maximization algorithm. IEEE transactions on medical imaging, 20 (1):45–57, 2001.

