## Supplementary Material for "Isolating the Sources of Pipeline-Variability in Group-Level Task-fMRI results"

### 8. Supplementary Methods and Results

#### 8.1. Consensus Analysis Method

For the ds000001 and ds000109 studies we performed the image-based meta-analysis originally proposed in Botvinik-Nezer et al., 2020 to quantify the evidence for brain activation across *all* analysis pipelines that had been applied to the data. While different meta-analytic approaches could be taken here (e.g. a random effects meta-analysis that penalizes for inter-pipeline variation), the key benefit of this particular 'consensus' analysis approach was that it accounted for the dependence between pipelines owing to the same underlying data and identical procedures implemented across parts of the analysis workflow. In particular, the consensus map is based on the mean of all pipelines' $z$-statistic maps, but is shifted and scaled by global factors so that the mean and variance are equal to the original image-wise means and variances averaged over all analysis workflows. Under the complete null of no signal across all voxels for every analysis workflow, the resulting consensus map can be expected to produce nominal standard normal $z$-scores. However, in the presence of signal these $z$-scores will reflect a consensus across the collection of results obtained from all individual analysis workflows.

The image-based meta-analysis method is as follows. Let $N$ be the total number of workflows through which the data has been analysed (for each of ds000001 and ds000109, we considered a total of 26 analysis pipelines), let $\mu$ be the (scalar) mean over space of each workflow's unthresholded $z$-statisic map, averaged across all pipelines, likewise let $\sigma^2$ be the spatial variance averaged over workflows, and let $\mathbf{Q}$ be the $N \times N$ correlation matrix, computed using all voxels in the statistical map. Then let $z_{ik}$ be the $z$-value for voxel $i$ and pipeline $k$, and $M_i$ the mean of those $N$ $z$-values at voxel $i$. The variance of $M_i$ is $\sigma^2 \mathbf{1}^\top \mathbf{Q} \mathbf{1} / N^2$ , where $\mathbf{1}$ is a $N$-vector of ones. We center and standardize $M_i$, and then rescale and shift to produce a meta-analytical $z$-map with mean $\mu$ and variance $\sigma^2$:

$$z_i = (M_i - \mu) \Big/ \sqrt{(\sigma^2 \mathbf{1}^\top \mathbf{Q} \mathbf{1} / N^2)} \times \sigma + \mu.$$

Two one-sided FDR voxelwise corrections were carried out at the 5% level to determine regions with significant positive and negative activation, using the two-stage linear step-up procedure (Benjamini et al., 2006).

#### 8.2. Overview of Results

Slice views of the thresholded statistic maps, correlation matrices, and Dice coefficient matrices are presented for all studies and pipelines in Supplementary Figures S7-S15.

Comparing the thresholded $t$-statistic maps presented in Supplementary Figures S7-S10 for ds000001, qualitative similarities are visible in the final brain regions of activation determined by most analysis workflows. For the majority of pipelines, positive activation was identified in the anterior cingulate, the anterior insula (bilateral), and the thalamus (bilateral), while negative activation was determined in the ventromedial prefrontal cortex and precuneus.

The main exception to this profile of activation was for the pipeline where preprocessing was carried out in fMRIPrep and FSL was used for the remainder of the analysis (pipeline **6AF/6SF** in Supplementary Figs. S7-S10), as well as the pipeline where the entire analysis was conducted in FSL (pipeline **7F** in Supplementary Figs. S7-S10). Specifically, the FSL pipeline with non-parametric inference at the group-level (pipeline **7F** in Supplementary Figs. S9 and S10) did not find any positive activations, and neither of the fMRIPrep/FSL pipelines identified activation in the anterior cingulate (pipelines **6AF/6SF** in Supplementary Figs. S7-S10). Further investigation of the unthresholded maps revealed that the lack of activation in the anterior cingulate here was likely due to the clusterwise inference performed for this study: while the other workflows obtained a single large activation cluster in the anterior cingulate, for the pipelines mentioned above this broke up into smaller, disconnected clusters, causing the activation to be 'thresholded out' after the FWE clusterwise correction. It is also notable that the FSL pipeline which used parametric inference (pipeline **7F** in Supplementary Figs. S7 and S8) found a small positive cluster in the visual cortex, which was not identified by any other pipelines. On the contrary, a *negative* effect was detected in the visual cortex for many of the AFNI/FSL hybrid pipelines shown in Supplementary Fig. S7 (pipelines **1A-5AF**). Finally, the activation clusters for pipelines that used AFNI's parametric group-level model (pipelines **1A** and **2AF** in Supplementary Fig. S7) and SPM's parametric group-level model (pipelines **1S** and **2SF** in Supplementary Fig. S8) generally have larger statistic values than the clusters for the remaining pipelines where FSL's parametric group-level model was used (pipelines **3AF/3SF-7F** in Supplementary Figs. S7 and S8).

Relative to ds000001, the thresholded $t$-statistic maps presented for ds000109 in Supplementary Figs. S11-S14 appear more qualitatively similar: across all workflows, strong positive effects were identified in the precuneus, frontal pole and superior frontal gyrus, bilateral superior occipital cortex and posterior temporal gyrus. On the other hand, greater variability arose between pipelines for delineating weaker effects. For example, there was disagreement across each collection of pipelines presented in Supplementary Figs. S11-S14 as to whether the posterior cingulate gyrus was positively activated or not (central activation cluster seen in the axial slice of the thresholded $t$-statistic map for some pipelines, but not others), alongside further discord as to whether *any* negative effects were present at all. Out of the 26 pipelines considered in total, 12 pipelines determined significant negative effects (8 using parametric group-level inference, 4 using nonparametric) while the other 14 did not. For the subset of results where negative effects were identified, clusters were often found in different brain regions with little-to-no overlap across workflows.

Similar to the ds000109 study, qualititative similarities can be seen in the main effects captured by all pipelines for the ds000120 dataset, while weaker effects were less robust. Activation was found by nearly all workflows in the occipital pole, bilateral occipital cortex, lingual gyrus and precuneus, as well as the supplementary motor cortex, middle frontal gyrus (bilateral) and thalamus (bilateral). However, there was greater variation in the areas where weaker effects were present, as seen by the different scatterings of smaller activation clusters in the axial slices of the thresholded $F$-statistic maps displayed in Suppelementary Fig. S15 (bottom row). Finally, it is notable that the activations in the occipital lobe for the pipelines applying SPM's group-level inference model (pipelines **3AS-7S**) seem to be less extended than the two pipelines (**1A** and **2AS**) which used AFNI's group-level model.

#### *8.3. Supplementary Results Figures*

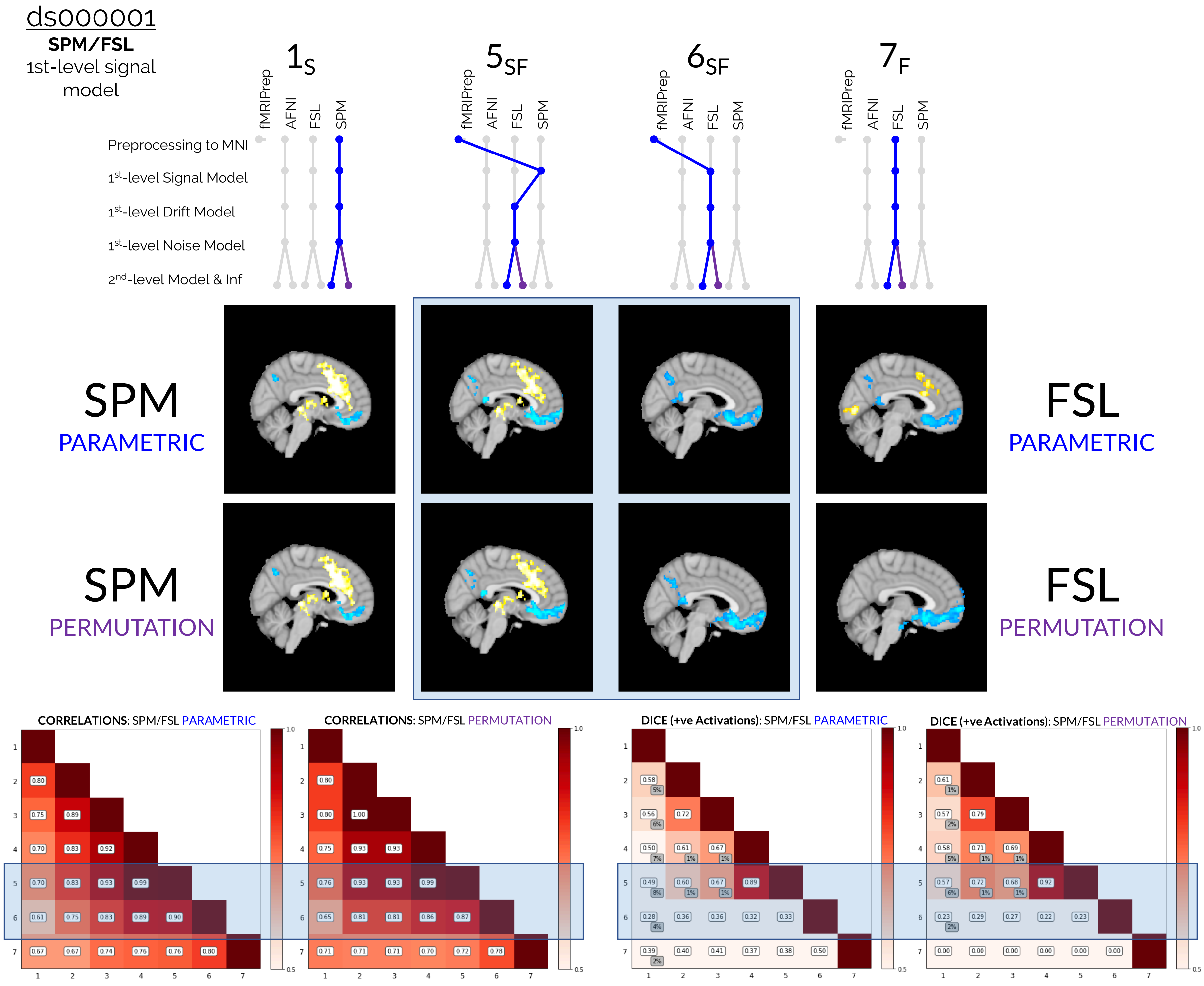
ds000001
SPM/FSL
1st-level signal model
1S
5SF
6SF
7F
fMRIPrep
AFNI
FSL
SPM
Preprocessing to MNI
1st-level Signal Model
1st-level Drift Model
1st-level Noise Model
2nd-level Model & Inf
SPM PARAMETRIC
SPM PERMUTATION
FSL PARAMETRIC
FSL PERMUTATION
CORRELATIONS: SPM/FSL PARAMETRIC
CORRELATIONS: SPM/FSL PERMUTATION
DICE (+ve Activations): SPM/FSL PARAMETRIC
DICE (+ve Activations): SPM/FSL PERMUTATION

Figure S1: Similar to Fig. 2, except this time focusing on the collection of results obtained from hybrid pipelines that implemented procedures from both SPM and FSL (rather than *AFNI* and FSL). Once again, the interchange of first-level signal model led to more extensive differences in the final results than any other individual processing stop, and similar to Fig. 2, this was largely due to the complete loss of positive activation in the thresholded maps that occured when SPM's first-level signal model (pipeline **5SF**) was interchanged with FSL's first-level signal model (pipeline **6SF**). Relative to the corresponding AFNI/FSL correlations presented in Fig. 2, the correlation values connected to pipeline **6SF** are improved here (bottom-left). This suggests that the overall differences in the activation profiles of the unthresholded maps for pipelines **5SF** and **6SF** were more subtle compared to the corresponding AFNI/FSL results, but that these differences were amplified after the FWE clusterwise correction was applied to obtain the thresholded maps.

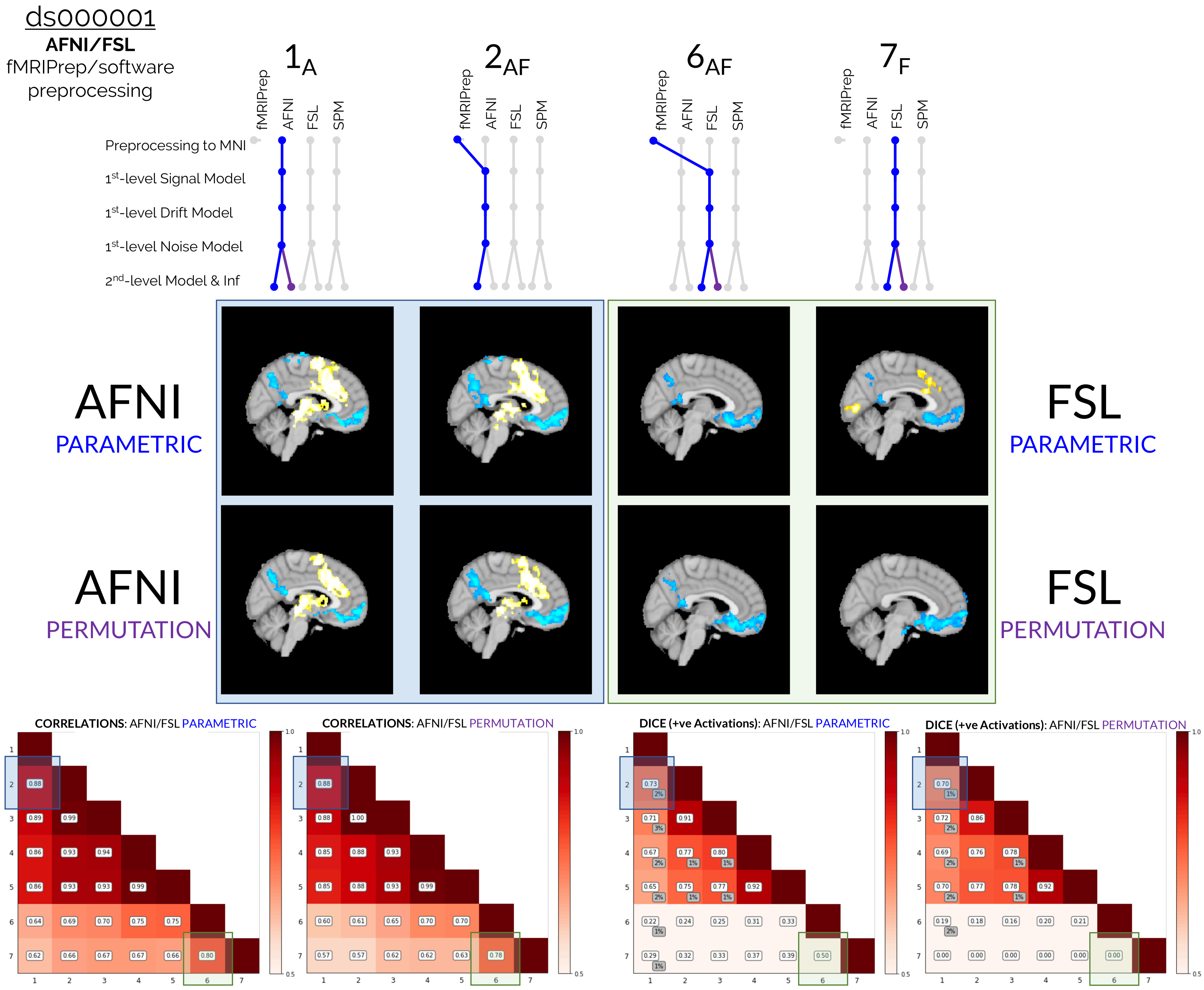
ds000001
AFNI/FSL
fMRIPrep/software preprocessing
1A
2AF
6AF
7F
fMRIPrep
AFNI
FSL
SPM
Preprocessing to MNI
1st-level Signal Model
1st-level Drift Model
1st-level Noise Model
2nd-level Model & Inf
AFNI PARAMETRIC
AFNI PERMUTATION
FSL PARAMETRIC
FSL PERMUTATION
CORRELATIONS: AFNI/FSL PARAMETRIC
0.88
0.89 0.99
0.86 0.93 0.94
0.86 0.93 0.93 0.99
0.64 0.69 0.70 0.75 0.75
0.62 0.66 0.67 0.67 0.66 0.80
CORRELATIONS: AFNI/FSL PERMUTATION
0.88
0.88 1.00
0.85 0.88 0.93
0.85 0.88 0.93 0.99
0.60 0.61 0.65 0.70 0.70
0.57 0.57 0.62 0.62 0.63 0.78
DICE (+ve Activations): AFNI/FSL PARAMETRIC
0.73 2%
0.71 3% 0.91
0.67 2% 0.77 1% 0.80 1%
0.65 2% 0.75 1% 0.77 1% 0.92
0.22 1% 0.24 0.25 0.31 0.33
0.29 1% 0.32 0.33 0.37 0.39 0.50
DICE (+ve Activations): AFNI/FSL PERMUTATION
0.70 1%
0.72 2% 0.86
0.69 2% 0.76 0.78 1%
0.70 2% 0.77 0.78 1% 0.92
0.19 2% 0.18 0.16 0.20 0.21
0.00 0.00 0.00 0.00 0.00 0.00
1 2 3 4 5 6 7
0.5
1.0

Figure S2: Comparisons of the group-level thresholded *t*-statistic maps, correlation values, and Dice coefficients obtained from reanalyses of the ds000001 dataset. Blue windows compare the two sets of results obtained from pipelines **1A** and **2AF**, which differed only as to whether preprocessing was carried out within AFNI or within fMRIPrep, respectively. Green windows compare pipelines **6AF** and **7F**, which differed only as to whether preprocessing was carried out within FSL or fMRIPrep. Qualitative and quantitative comparisons displayed here show a high degree of similarity between the two sets of results where either AFNI or fMRIPrep preprocessing was used, while greater differences can be seen for the two pipelines where FSL's preprocessing workflow was interchanged with fMRIPrep. In particular, the slice views of the thresholded *t*-statistic maps for pipelines **1A** and **2AF** look strikingly similar (middle, blue window), while disagreement can be seen in terms of the brain regions that were positively activated for the parametric results obtained with pipelines **6AF** and **7F** (middle, green window). Alongside this, the correlation and Dice values for the pairwise comparisons of the **1A** and **2AF** results (blue windows, bottom-left and bottom-right) were better than the corresponding values obtained for **6AF** and **7F** regardless of whether parametric or nonparametric inference was performed.

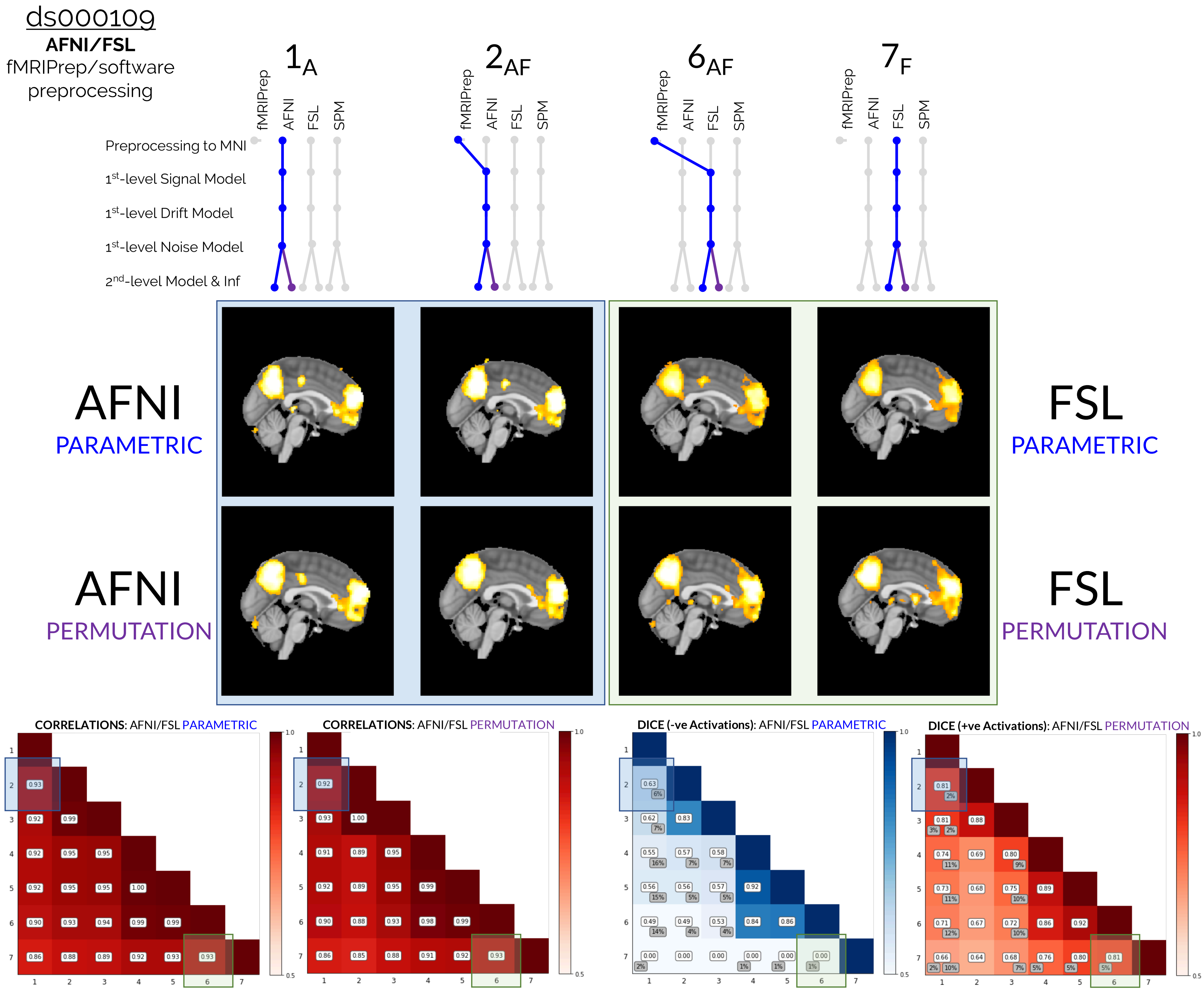
ds000109
AFNI/FSL
fMRIPrep/software preprocessing
$1_A$
$2_{AF}$
$6_{AF}$
$7_F$
fMRIPrep
AFNI
FSL
SPM
Preprocessing to MNI
1st-level Signal Model
1st-level Drift Model
1st-level Noise Model
2nd-level Model & Inf
AFNI
PARAMETRIC
AFNI
PERMUTATION
FSL
PARAMETRIC
FSL
PERMUTATION
CORRELATIONS: AFNI/FSL PARAMETRIC
CORRELATIONS: AFNI/FSL PERMUTATION
DICE (-ve Activations): AFNI/FSL PARAMETRIC
DICE (+ve Activations): AFNI/FSL PERMUTATION

Figure S3: Similar to Fig. S2, except this time focusing on the corresponding pipeline results for reanalyses of the ds000109 dataset. While greater similarity can be seen on-the-whole for the ds000109 results compared to ds000001, it is notable that there was still disagreement between pipelines **6AF** and **7F** in terms of the negatively activated brain regions for the parametric inference results: while the thresholdd results for pipeline **6AF** (that used fMRIPrep's preprocessing workflow) determined two clusters of negative activation in the inferior temporal gyrus (bilateral), pipeline **7F** (identical to **6AF** except that FSL's preprocessing was used) didn't determine *any* negative activation. Consequently, the Dice coefficient for comparisons of these two pipelines is zero (green window in the blue negative activation Dice matrix at bottom).

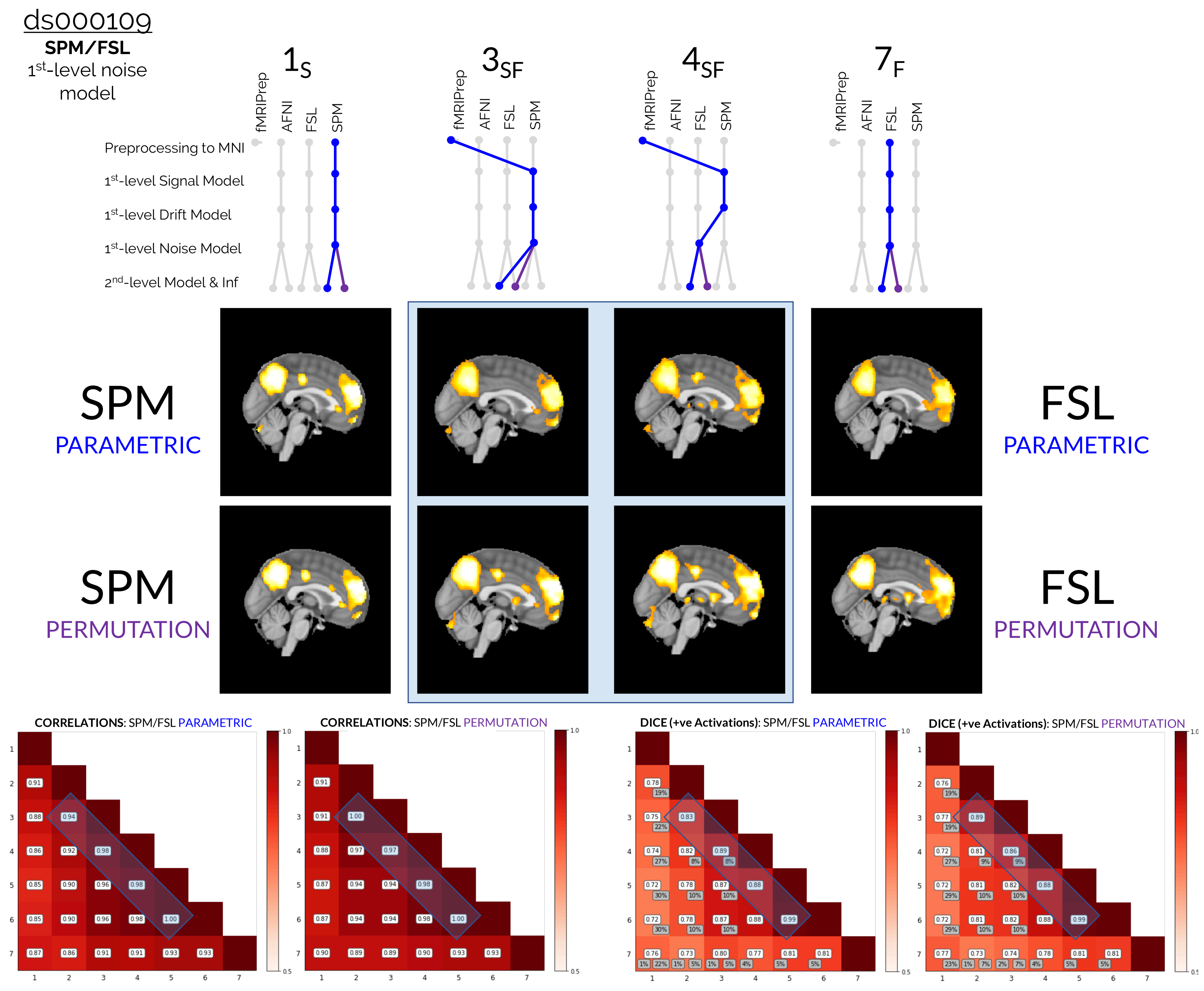

ds000109
SPM/FSL
1st-level noise model
1S
3SF
4SF
7F
fMRIPrep
AFNI
FSL
SPM
Preprocessing to MNI
1st-level Signal Model
1st-level Drift Model
1st-level Noise Model
2nd-level Model & Inf
SPM PARAMETRIC
SPM PERMUTATION
FSL PARAMETRIC
FSL PERMUTATION
CORRELATIONS: SPM/FSL PARAMETRIC
CORRELATIONS: SPM/FSL PERMUTATION
DICE (+ve Activations): SPM/FSL PARAMETRIC
DICE (+ve Activations): SPM/FSL PERMUTATION

Figure S4: Similar to Fig. 3, except this time focusing on the collection of results obtained from hybrid pipelines that implemented procedures from both SPM and FSL (rather than *AFNI* and FSL). Once again, setting preprocessing aside, the interchange of the first-level noise model between pipelines **3SF** and **4SF** led to more extensive differences in the results than any other modelling procedure. This is highlighted in the blue windows on the off-diagonals of the correlation and Dice matrices at the bottom of the figure, where the values for pipelines **3SF** and **4SF** can be seen to be lower than the corresponding values for other pairs of adjacent pipelines in most cases. Similarly to Fig. 3, the thresholded $t$-statistic map for pipeline **4SF** (that used FSL's first-level noise model) determined slightly more smaller activation clusters than the corresponding set of results for pipeline **3SF** (that used SPM's noise model).

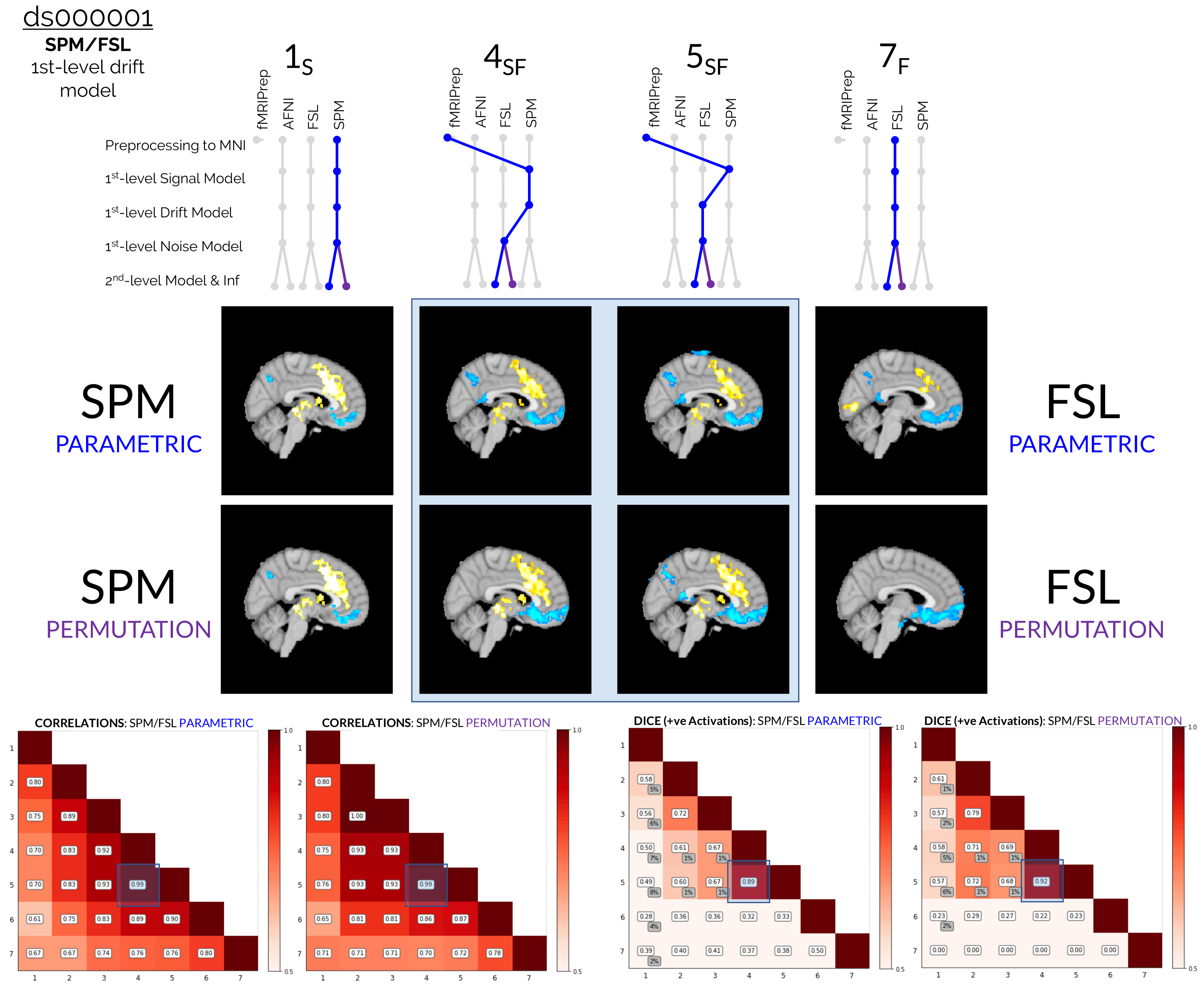
ds000001
SPM/FSL
1st-level drift model
$1_S$
$4_{SF}$
$5_{SF}$
$7_F$
fMRIPrep
AFNI
FSL
SPM
Preprocessing to MNI
1st-level Signal Model
1st-level Drift Model
1st-level Noise Model
2nd-level Model & Inf
SPM
PARAMETRIC
SPM
PERMUTATION
FSL
PARAMETRIC
FSL
PERMUTATION
CORRELATIONS: SPM/FSL PARAMETRIC
CORRELATIONS: SPM/FSL PERMUTATION
DICE (+ve Activations): SPM/FSL PARAMETRIC
DICE (+ve Activations): SPM/FSL PERMUTATION

Figure S5: Comparisons of the group-level thresholded *t*-statistic maps, correlation values, and Dice coefficients obtained from reanalyses of the ds000001 dataset, focusing on the collection of results obtained from hybrid pipelines that implemented procedures from both SPM and FSL. The two sets of results given by pipelines **4SF** and **5SF** are displayed, which differed only as to whether SPM's or FSL's first-level drift model was used. Overall, the change of drift model had minimal impact on the final results; pairwise comparisons show that the unthresholded maps for pipelines **4SF** and **5SF** were almost perfectly correlated for both the parametric and nonparametric inference cases (bottom-left, blue windows). The Dice values are marginally worse (bottom-right, blue window) – close to 90% – due to slightly more negative activation determined by pipeline **5SF** (that used FSL's first-level drift model) as seen in the thresholded *t*-statistic maps (middle, blue window). Nevertheless, the correlations and Dice comparisons for pipelines **4SF** and **5SF** are the best of all pairs of adjacent pipelines.

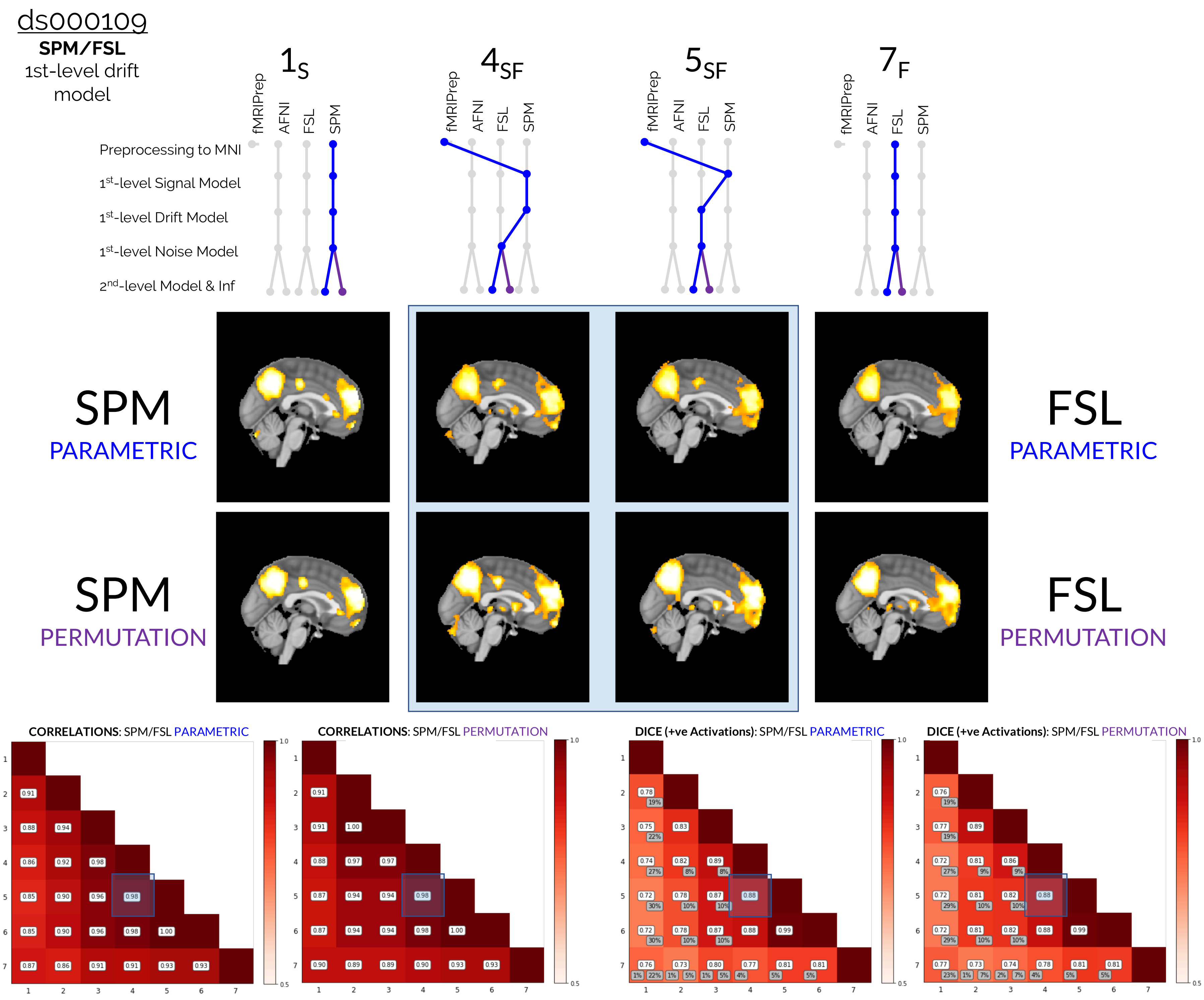

ds000109
SPM/FSL
1st-level drift model
1S
4SF
5SF
7F
fMRIPrep
AFNI
FSL
SPM
Preprocessing to MNI
1st-level Signal Model
1st-level Drift Model
1st-level Noise Model
2nd-level Model & Inf
SPM PARAMETRIC
SPM PERMUTATION
FSL PARAMETRIC
FSL PERMUTATION
CORRELATIONS: SPM/FSL PARAMETRIC
CORRELATIONS: SPM/FSL PERMUTATION
DICE (+ve Activations): SPM/FSL PARAMETRIC
DICE (+ve Activations): SPM/FSL PERMUTATION

Figure S6: Similar to Fig. S5, except this time focusing on the corresponding pipeline results for reanalyses of the ds000109 dataset. Once again, the correlations (blue windows, bottom-left) and Dice values (blue windows, bottom-right) for pairwise comparisons of pipelines **4SF** and **5SF** were some of the best of any, indicating that the interchange of drift model between SPM and FSL minimally impacted the final group-level results. In the thresholded *t*-statistic images (blue winodw, middle), it can be seen that the change of drift model between the two software packages only led to slight changes in the locations of some of the smaller activation clusters.

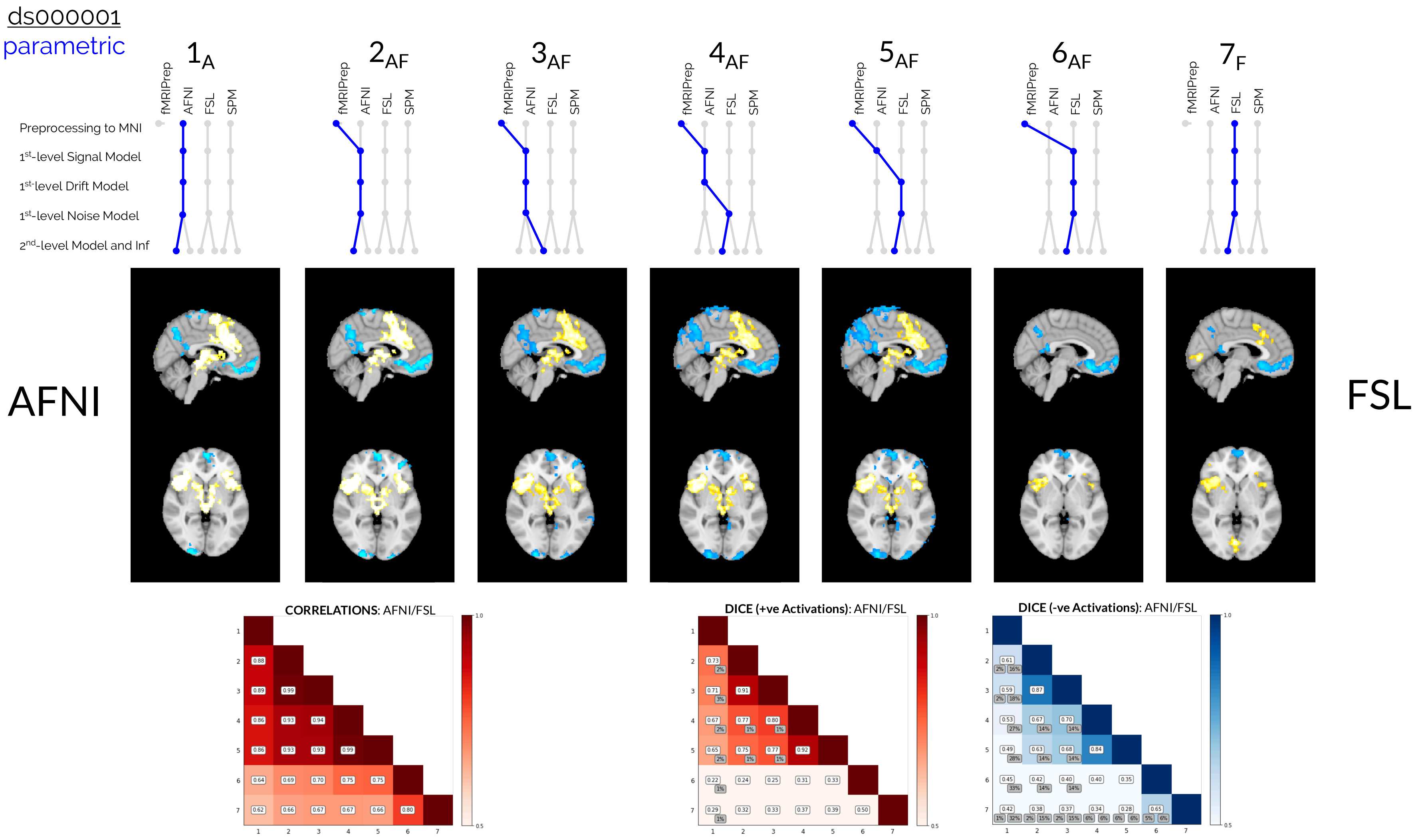


Figure S7: **ds000001 AFNI/FSL Pipelines (Parametric Results).** Comparisons of the group-level thresholded *t*-statistic maps, correlation values, and Dice coefficients obtained from reanalyses of the ds000001 dataset. The collection of all parametric inference results obtained from hybrid pipelines that implemented procedures from both AFNI and FSL are presented.

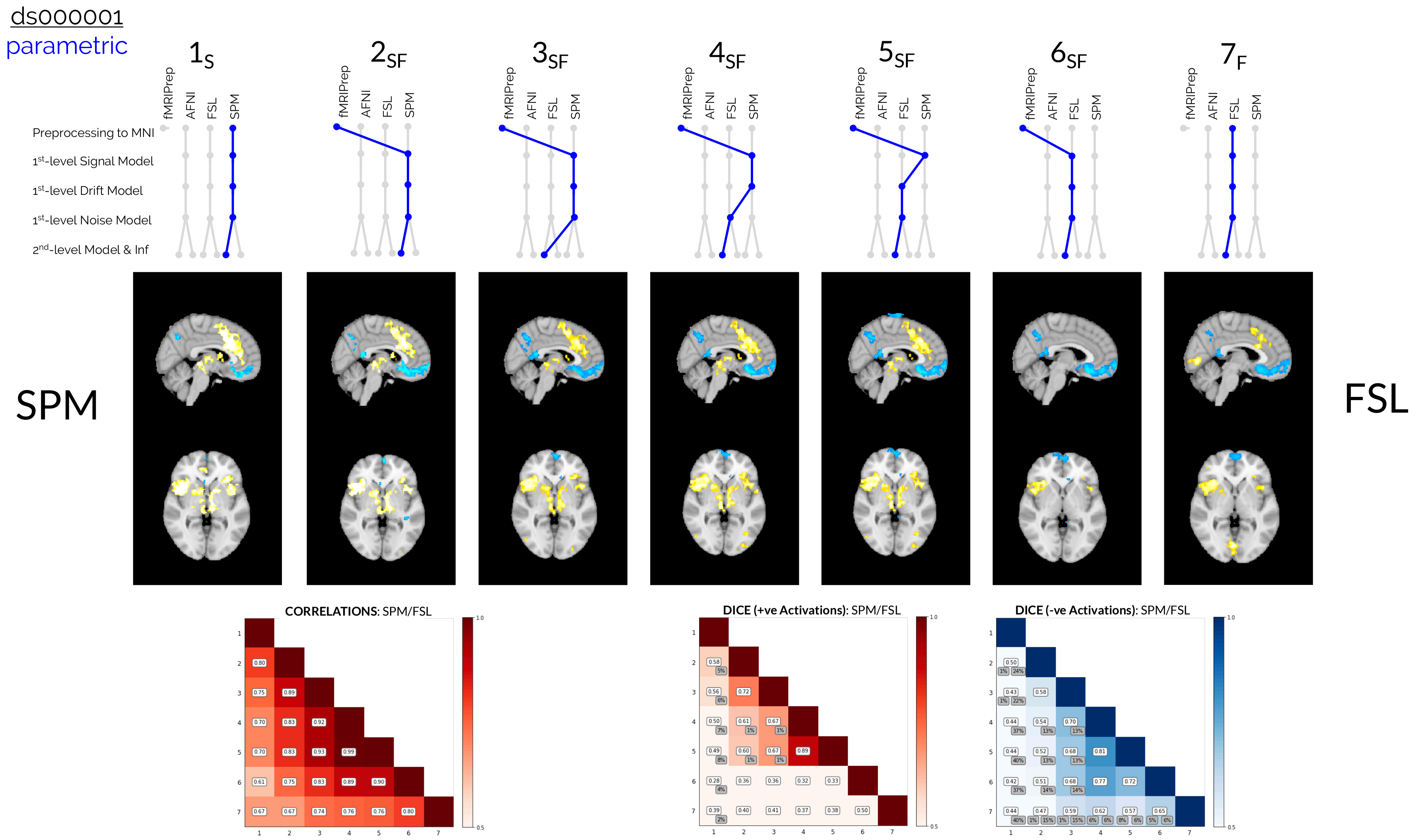


Figure S8: **ds000001 SPM/FSL Pipelines (Parametric Results).** Comparisons of the group-level thresholded *t*-statistic maps, correlation values, and Dice coefficients obtained from reanalyses of the ds000001 dataset. The collection of all results obtained from hybrid pipelines that implemented procedures from both AFNI and FSL are presented.

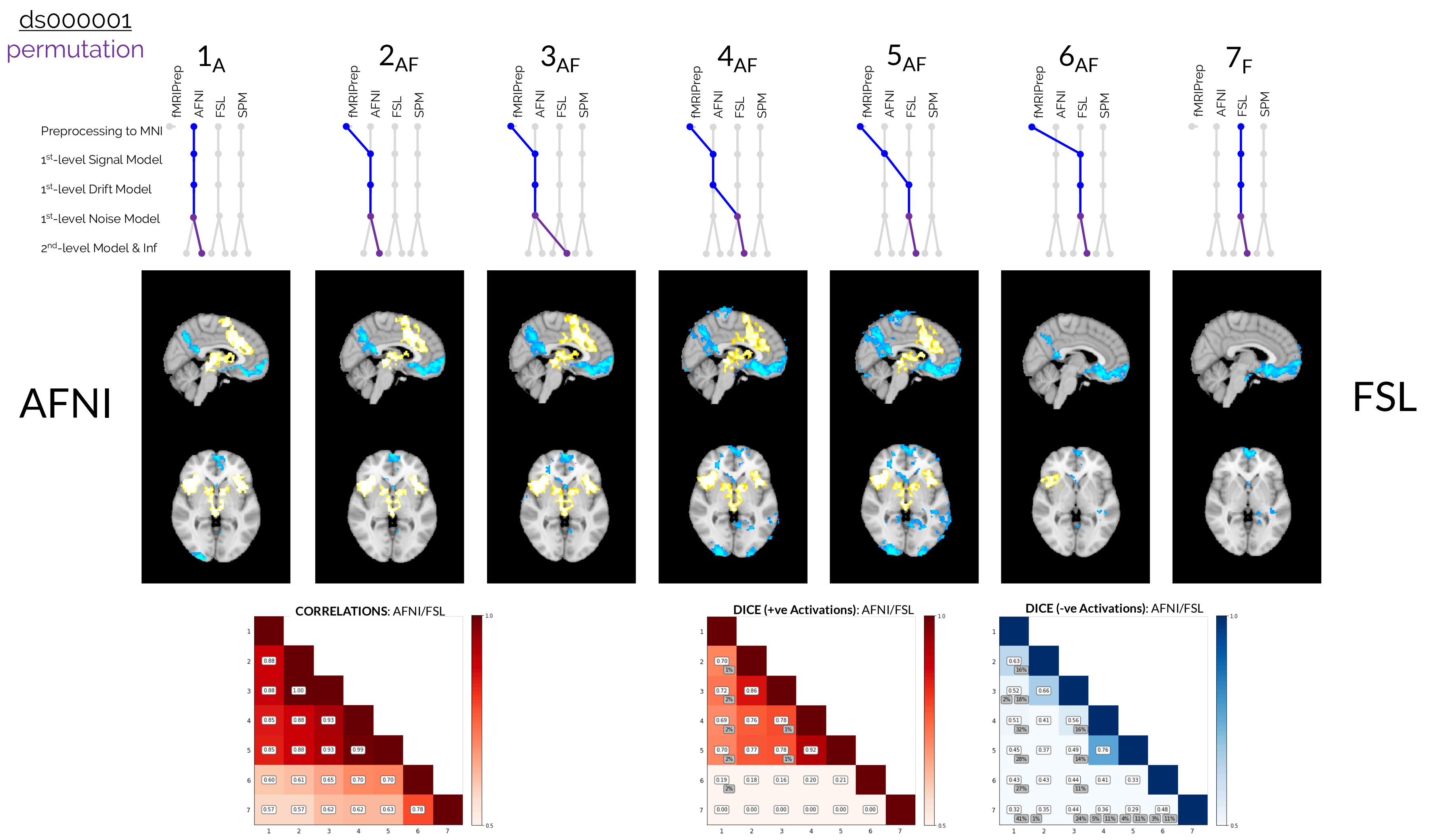


Figure S9: **ds000001 AFNI/FSL Pipelines (Nonparametric Results).** Comparisons of the group-level thresholded *t*-statistic maps, correlation values, and Dice coefficients obtained from reanalyses of the ds000001 dataset. The collection of all nonparametric inference (permutation test) results obtained from hybrid pipelines that implemented procedures from both AFNI and FSL are presented.

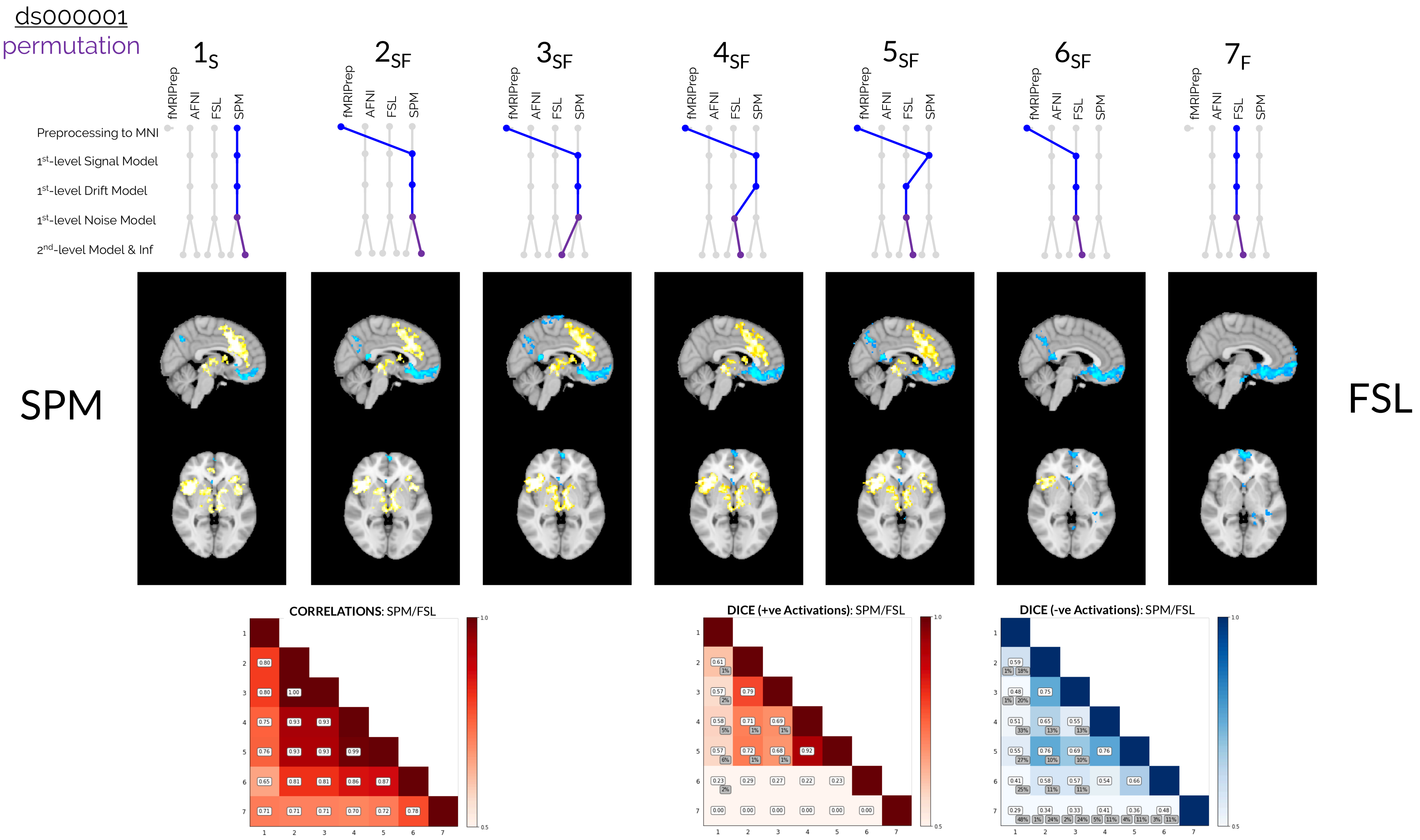


Figure S10: **ds000001 SPM/FSL Pipelines (Nonparametric Results).** Comparisons of the group-level thresholded $t$-statistic maps, correlation values, and Dice coefficients obtained from reanalyses of the ds000001 dataset. The collection of all nonparametric inference (permutation test) results obtained from hybrid pipelines that implemented procedures from both SPM and FSL are presented.

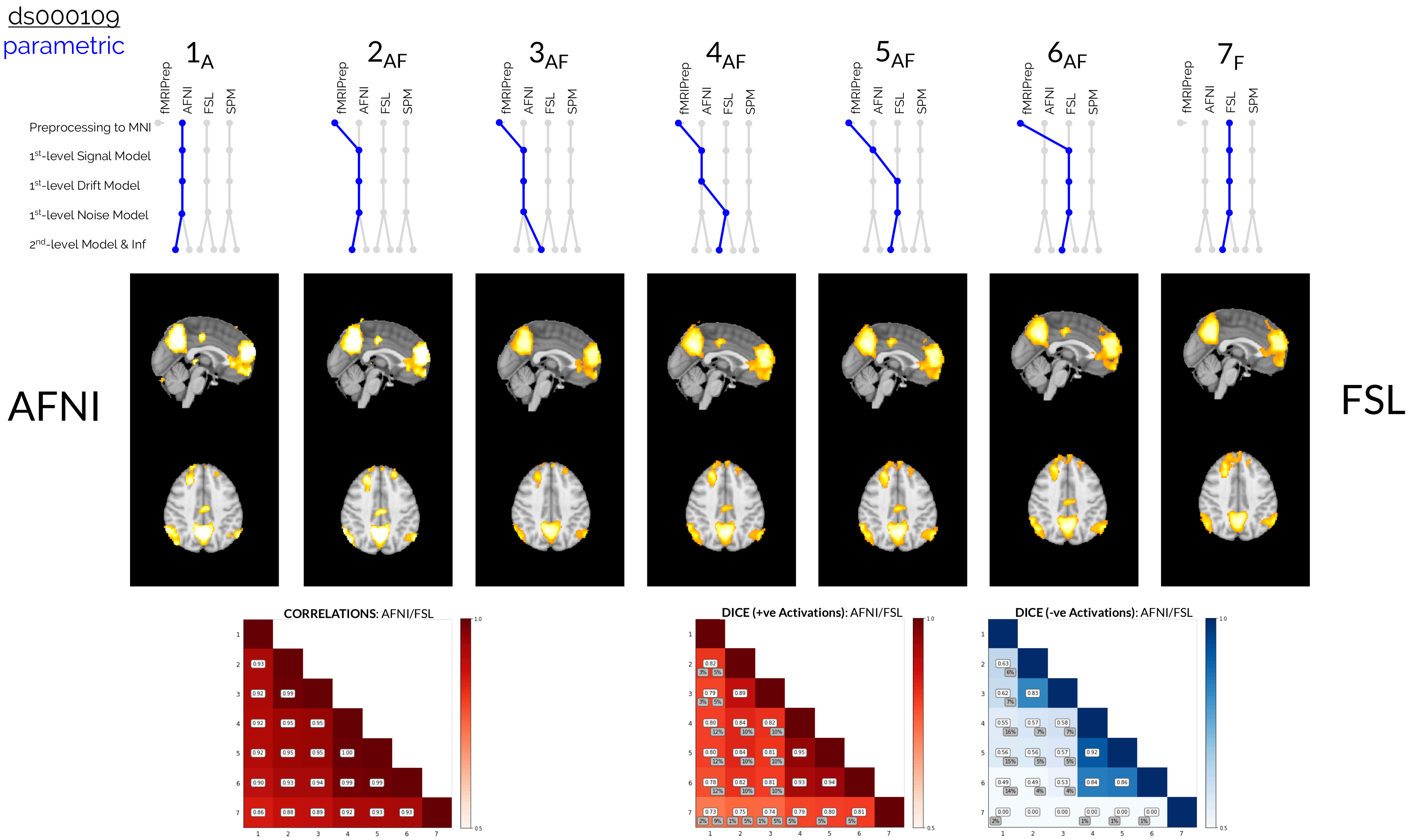


Figure S11: **ds000109 AFNI/FSL Pipelines (Parametric Results).** Comparisons of the group-level thresholded $t$-statistic maps, correlation values, and Dice coefficients obtained from reanalyses of the ds000109 dataset. The collection of all parametric inference results obtained from hybrid pipelines that implemented procedures from both AFNI and FSL are presented.

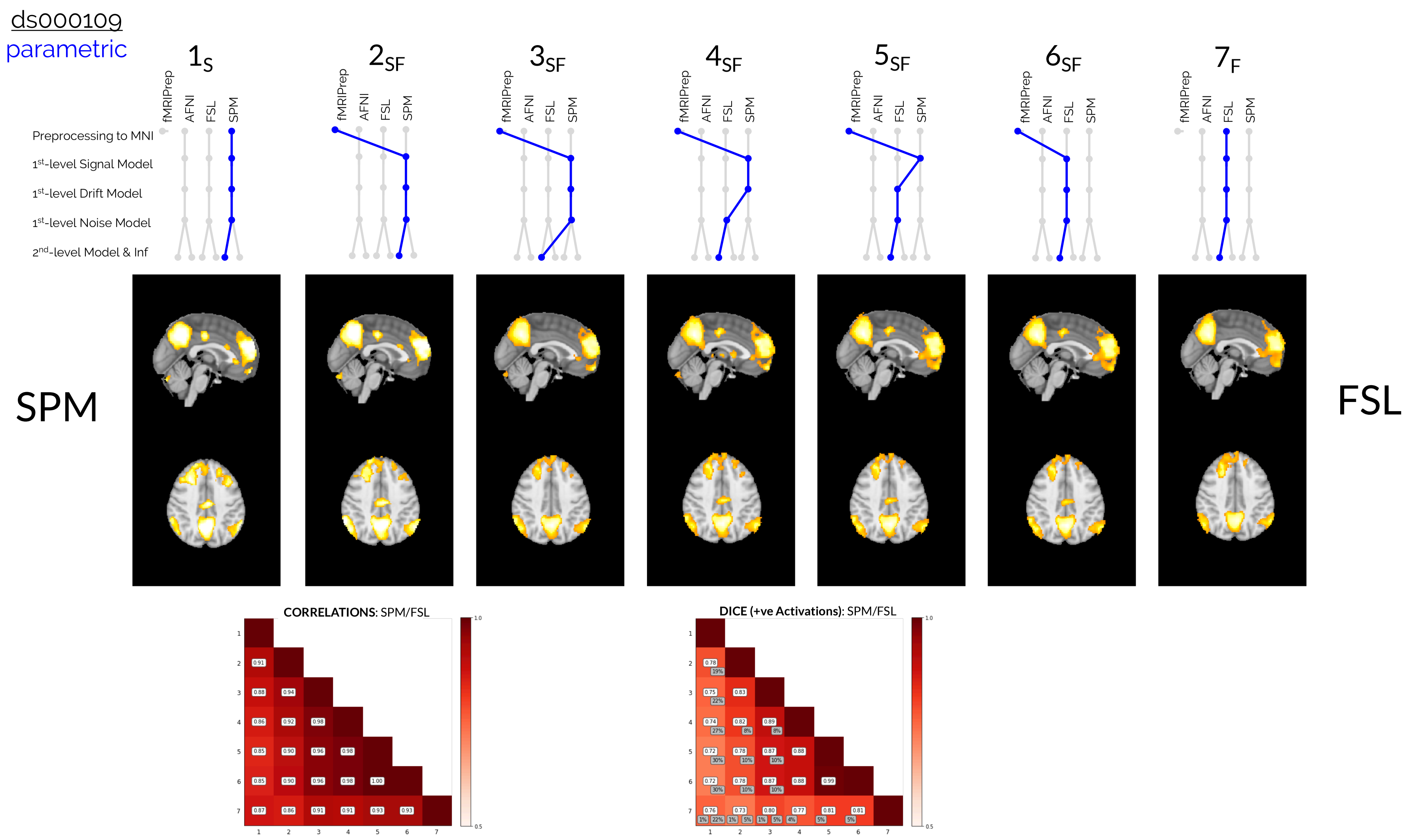


Figure S12: **ds000109 AFNI/FSL Pipelines (Nonparametric Results).** Comparisons of the group-level thresholded *t*-statistic maps, correlation values, and Dice coefficients obtained from reanalyses of the ds000109 dataset. The collection of all nonparametric inference (permutation test) results obtained from hybrid pipelines that implemented procedures from both AFNI and FSL are presented.

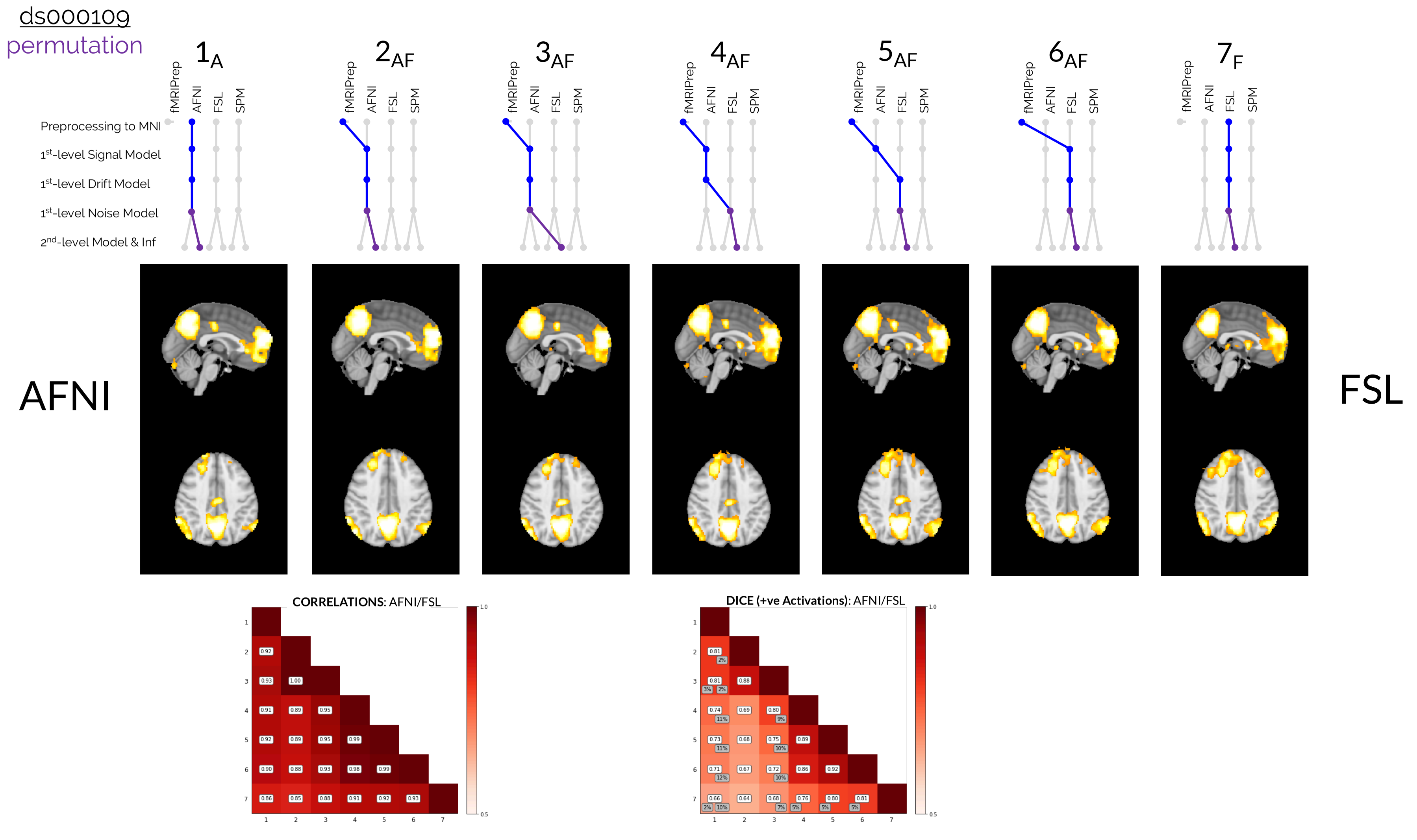


Figure S13: **ds000109 AFNI/FSL Pipelines (Nonparametric Results).** Comparisons of the group-level thresholded *t*-statistic maps, correlation values, and Dice coefficients obtained from reanalyses of the ds000109 dataset. The collection of all nonparametric inference (permutation test) results obtained from hybrid pipelines that implemented procedures from both AFNI and FSL are presented.

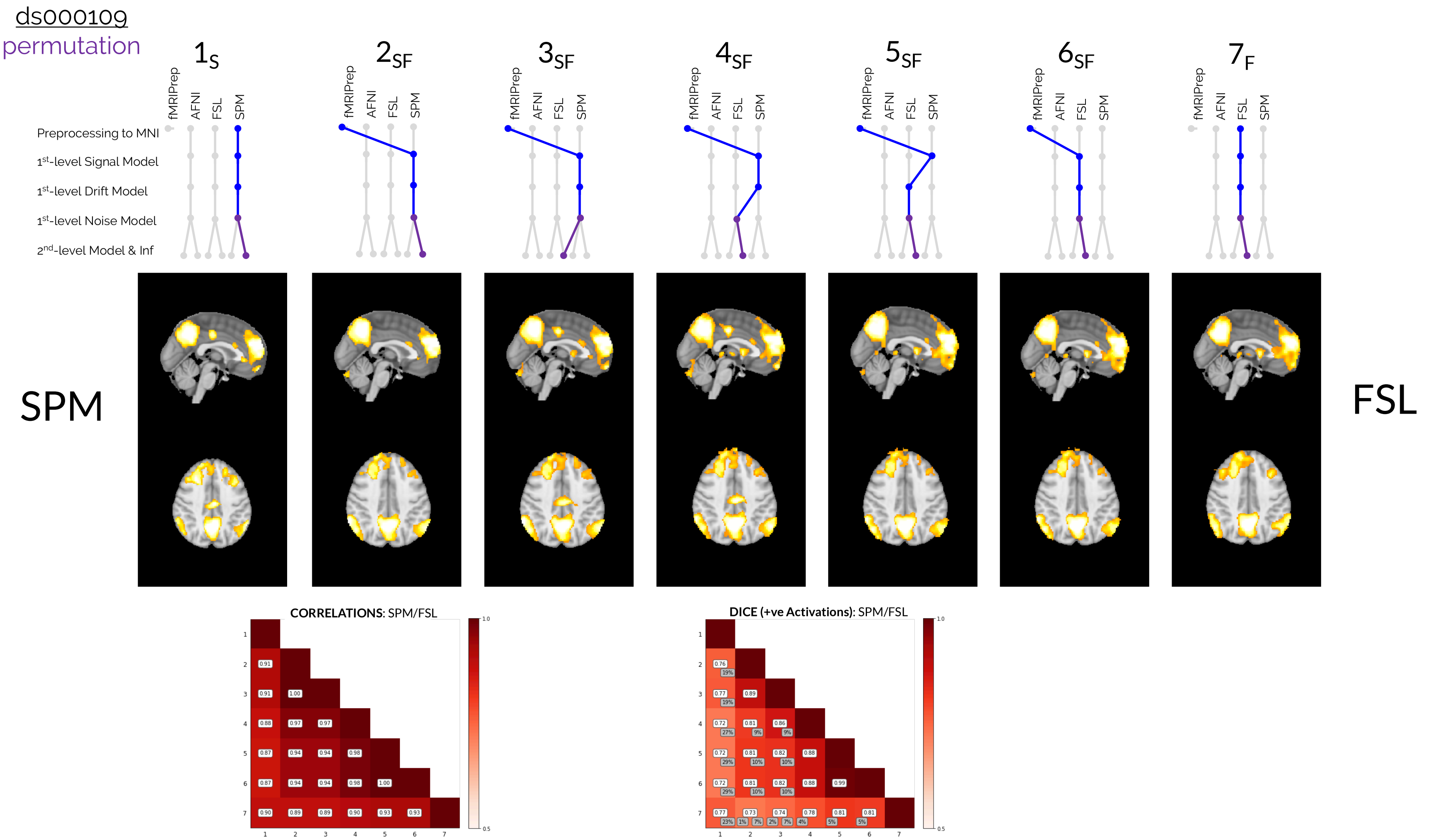


Figure S14: **ds000109 SPM/FSL Pipelines (Nonparametric Results).** Comparisons of the group-level thresholded $t$-statistic maps, correlation values, and Dice coefficients obtained from reanalyses of the ds000109 dataset. The collection of all nonparametric inference (permutation test) results obtained from hybrid pipelines that implemented procedures from both SPM and FSL are presented.

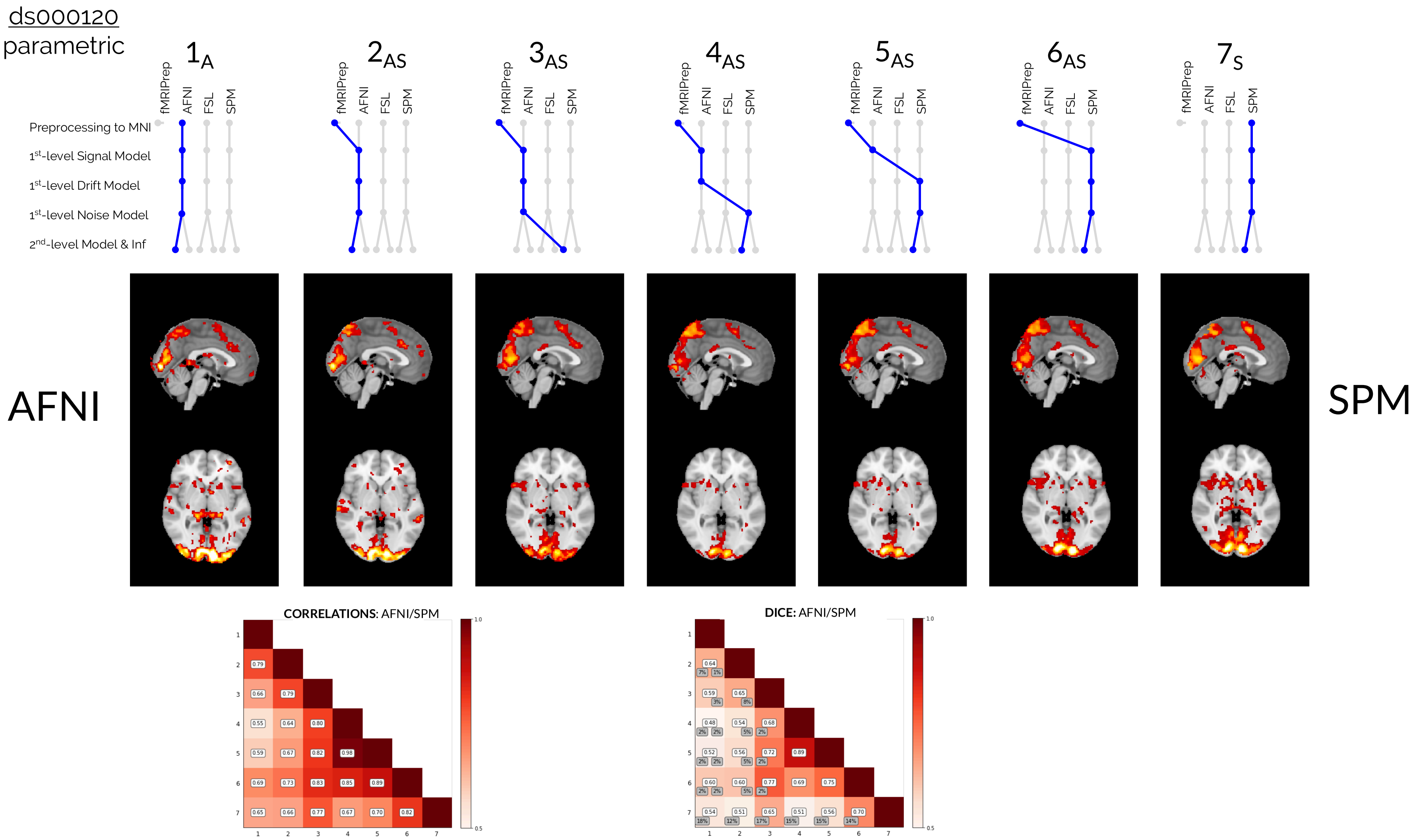


Figure S15: **ds000120 AFNI/SPM Pipelines (Parametric Results).** Comparisons of the group-level thresholded $F$-statistic maps, correlation values, and Dice coefficients obtained from reanalyses of the ds000120 dataset. The collection of all parametric inference results obtained from hybrid pipelines that implemented procedures from both AFNI and SPM are presented.
